# Mechanosensitive ion channel PIEZO1 enhances endometrial decidualization through BECN1-dependent autophagy

**DOI:** 10.1101/2025.08.05.668790

**Authors:** Jin-Wen Kang, Yao Wu, Yu Zhang, Hui-Xia Li, Guang-Ya Li, Yao-Feng Yang, Xiao-Qing Huang, Jia-Ying Yu, Chen Liang, Rui Zhang, Xiao-Zheng Liu, Shan-Shan Song, Ying-Nan Liu, Aftab Shaukat, Yong Song, Samantha Hrbek, John Lydon, Bin Guo, Hong-Lu Diao, Zeng-Ming Yang, Asgerally Fazleabas, Ren-Wei Su

## Abstract

The mechanosensitive ion channel PIEZO1 plays critical roles in physiological and pathological processes in response to various types of mechanical forces, including shear stress, stretch, and extracellular matrix (ECM) stiffness. Decidualization is crucial for a successful pregnancy, characterized by the differentiation of fibroblastic endometrial stromal cells into round, secretory decidual cells, along with the rapid remodeling of the ECM. Herein, we report that PIEZO1 plays a crucial role in enhancing decidualization in response to extracellular matrix (ECM) stiffness and cell contraction. Uterine-specific knockout of *Piezo1* using *Pgr-Cre* in mice results in subfertility due to decidualization impairment in mid-late pregnancy. Silencing of *PIEZO1* in human endometrial stromal cells also results in impaired decidualization. Treatment with the PIEZO1 agonist Yoda1 enhances decidualization in both *in vivo* and *in vitro* models. Stromal cells growing on ECM with 25 kPa stiffness display a better decidualization response than cells seeded on softer 2 kPa surface or harder surface of the regulator petri dish, and this difference is abolished by null of Piezo1. Consistent with PIEZO1 as a Ca^2+^ modulator, blocking of intracellular Ca^2+^ or pCaMKII significantly inhibits Yoda1-enhanced decidualization. Further investigation reveals that BECN1-dependent autophagy acts as the downstream of PIEZO1. Silencing of *Beclin1* abolishes Yoda1-induced decidualization, while Tat-BECN1 fully rescues impaired decidualization caused by the lack of PIEZO1. Finally, the lower expression of *PIEZO1* is associated with impaired decidualization in the endometrium of endometriotic baboons. In conclusion, we have uncovered a novel mechanism of decidualization that is regulated by PIEZO1-mediated mechanotransduction, providing further insight into decidualization studies.

**Graphical abstract:** 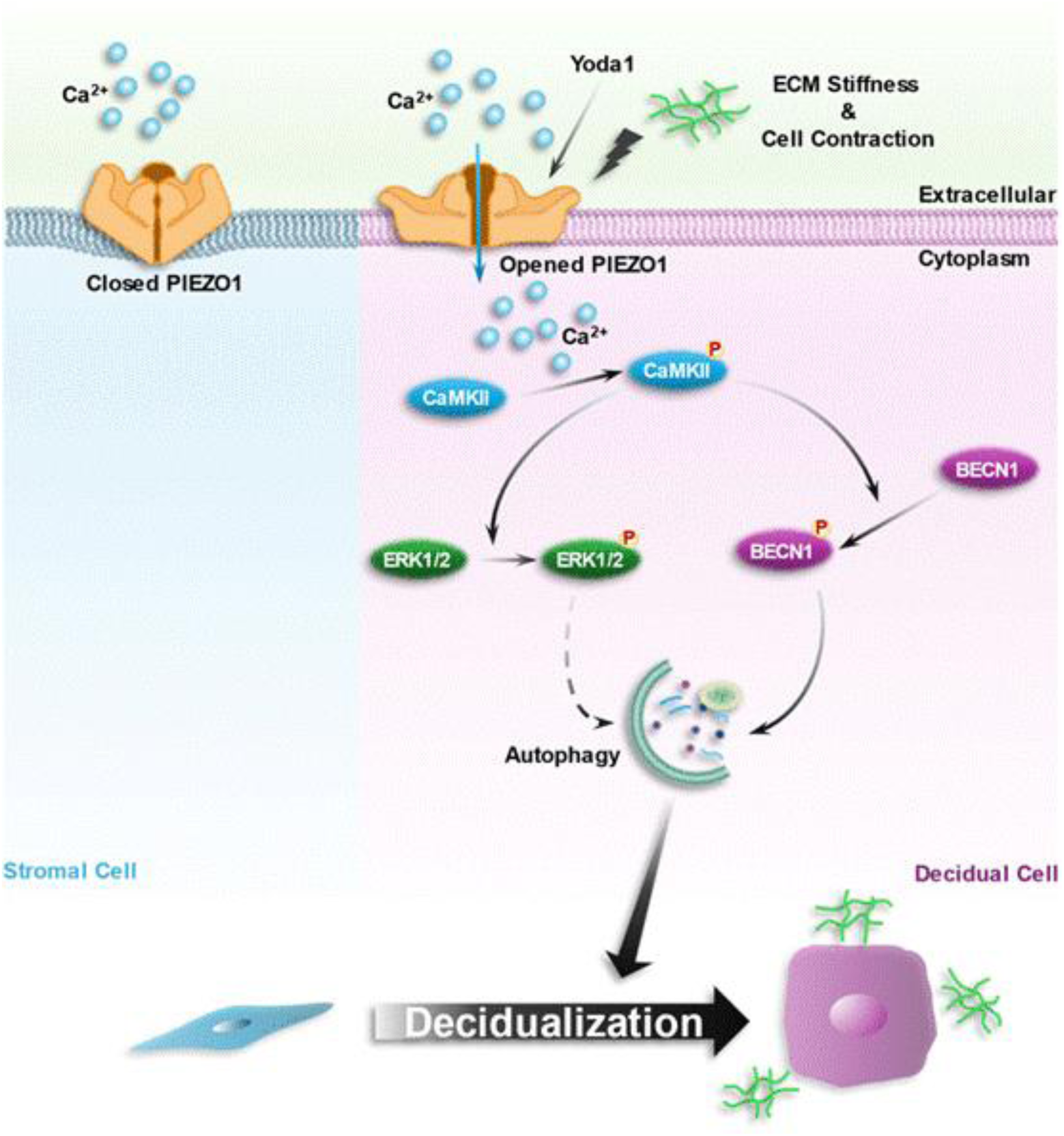

## Introduction

Decidualization is a highly regulated and essential process during pregnancy in primates and rodents, which defines as the trans-differentiation of fibroblastic endometrial stromal cells into specialized epithelial-like secretory decidual cells, accompany with rapid extracellular matrix (ECM) remodeling [1]. Disrupted decidualization is associated with several pregnancy-related complications, such as recurrent implantation failure, recurrent pregnancy loss, preeclampsia, preterm birth, and infertility/subfertility caused by endometriosis. Therefore, decidualization is considered “the primary driver of pregnancy health” [2–6]. Over the past decades, numerous factors, including two dominant ovarian hormones, estrogen, and progesterone, growth factors, cytokines, endocrine factors, embryonic signals, kinases and transcriptional factors, and the implanting embryo itself, have been identified as key regulators of decidualization [1, 7, 8]. In addition to these factors, the morphological alterations of endometrial stromal cells during decidualization and the changes in local stiffness and tissue tension caused by ECM remodeling may also be one of the regulators. Although mechanical forces are increasingly recognized as significant contributors during pregnancy, whether or not endometrial stromal cells integrate mechanical force signals from their *in vivo* microenvironment and the underlying mechanism of mechanical forces driving decidualization is still not understood (reviewed in [9]).

Cells convert mechanical stimulations, like hydrostatic pressure (HP), fluid shear stress (FSS), extracellular matrix (ECM) stiffness, and tensile force (TF) into cellular signals, known as mechanotransduction, which represents an indispensable biological function with evolutionary conservation [10]. Mechanically activated channels transfer force sensitivity to cells and organisms by allowing the passage of ions across the membrane in response to a mechanical stimulus, which triggers the electrochemical signals intercellularly [11]. In 2010, the mechanosensitive ion channel proteins PIEZO1 and PIEZO2 were identified as crucial mechanotransducers responsible for converting mechanical force into electrochemical signals [12]. PIEZO channels exhibit remarkable sensitivity to mechanical force, through interactions with lipid membranes, the cytoskeleton, or ECM [13, 14]. Over the past decade, PIEZO1 has emerged as a mechanosensitive ion channel with critical roles in numerous physiological and pathological processes, including cardiovascular [15], bone homeostasis [16], pressure-induced pancreatitis [17], lung functions [18], immune functions [19–21], and intestinal stem cell fate decision [22].

Previous studies have shown that during pregnancy, the distending forces from the growing deciduoma, fetus, and placenta progressively enhance tissue mechanics[23]. Using atomic force microscopy, researchers have observed significantly higher elastic modulus in *ex vivo* tissue samples obtained from the decidua basalis, the endometrial site of placental invasion, when compared to endometrial tissue samples from non-pregnant women (1250 Pa vs. 250 Pa), suggesting that invading extravillous may enhance decidual stiffness by remodeling vasculature and ECM [24]. Furthermore, studies have reported that the cycling mechanical stretch (occurring twice per minute) triggers the up-regulation of *IGFBP1*, a well-known decidualization marker, in an *in vitro* cultured decidualization model [25]. However, when the stretch frequency is increased (six times per minute), it adversely affects decidualization [26]. Another study indicates that stretch force leads to elevated cAMP levels and subsequent downstream signaling which are critical for decidualization, as well as the expression of αSMA, an early decidualization marker [27]. These investigations strongly demonstrate that endometrial stromal cell decidualization may be regulated by mechanical forces from its *in vivo* microenvironment. However, the specific molecular mechanism involved in the regulation of mechanical force-induced decidualization remains poorly understood.

Autophagy is an evolutionarily conserved biological process in eukaryotic cells that plays an important role in regulating cell metabolic homeostasis [28]. Three critical protein complexes contribute to the initiation and nucleation of autophagy: the ULK1 initiation complex which is negatively regulated by mTOR signaling, the PI3K III nucleation complex, and the PI3P-binding complex [29]. Autophagy has been shown to be enhanced during the decidualization process[30]. Inhibiting autophagy impairs decidualization in both mouse models and human cells[30–34]. Clinically, autophagy is decreased in endometrial stromal cells of patients with recurrent miscarriage and is associated with impaired decidualization capability [35]. On the other hand, mechanical cues, such as microgravity, stretch, and ECM stiffness are reported to affect autophagy [36–38]. However, whether autophagy acts as a mediator of mechanical cues, especially PIEZO1-mediated mechanotransduction, in regulating decidualization is still largely unknown.

In this study, we identified that PIEZO1 is functionally expressed during embryo implantation and decidualization. Uterine-specific knockout of *Piezo1* in mice resulted in significantly reduced fertility due to impaired decidualization during mid-late pregnancy compared to control mice. Furthermore, we demonstrated that specific mechanical cues, cell contraction, and ECM stiffness, regulate decidualization in both mice and human cells through PIEZO1. Detailed mechanistic analyses revealed dual downstream signaling pathways: Ca^2+^-CaMKII-ERK and Ca^2+^-CaMKII-BECN1-autophagy. Importantly, we uncovered that PIEZO1 may serve as a potential therapeutic target in impaired decidualization and endometriosis, offering new insights into its pathophysiology and treatment strategies.

## Results

### Spatiotemporal expression of functional PIEZO1 in the endometrium

We initially examined the expression pattern of PIEZO1 protein in the endometrium during early pregnancy using *Piezo1^tdTomato^* mice, which carry a *Piezo1* gene fused with a sequence encoding tdTomato. Application of both anti-RFP and anti-PIEZO1 antibodies immunostaining revealed that the robust spatiotemporal expression of PIEZO1-tdTomato fused protein in the decidualized zones from 4.5 to 7.5 days post-coitum (dpc) (Fig. 1A; Fig. S1A). In an artificial decidualization mouse model, PIEZO1 expression was significantly up-regulated in the decidualized horn compared to the non-stimulated control horn (Fig. 1B-D). Furthermore, we wondered if the expression of PIEZO1 was correlated with endometriosis, a gynecological disease that negatively affects decidualization [4]. We used a well-established non-human primate endometriotic model in which endometriosis was artificially induced by the intraperitoneal inoculation of menstrual tissue in baboons [39]. The results showed that the mRNA levels of *PIEZO1* were significantly down-regulated in the eutopic endometrium of baboons 15 months after the induction of disease compared to the disease-free endometrium from the same animal prior to the induction of endometriosis (Fig. 1E), which correlated with the impaired decidualization associated with this disease [4]. Moreover, applying mechanical stretching or treating with Yoda1, an agonist of PIEZO1 protein, to *ex vivo* cultured endometrial tissue up-regulated *Piezo1* mRNA levels within 5 minutes, suggesting a rapid self-regulation of *Piezo1* expression in response to mechanical stimulation (Fig1. F&G).

**Figure 1.**
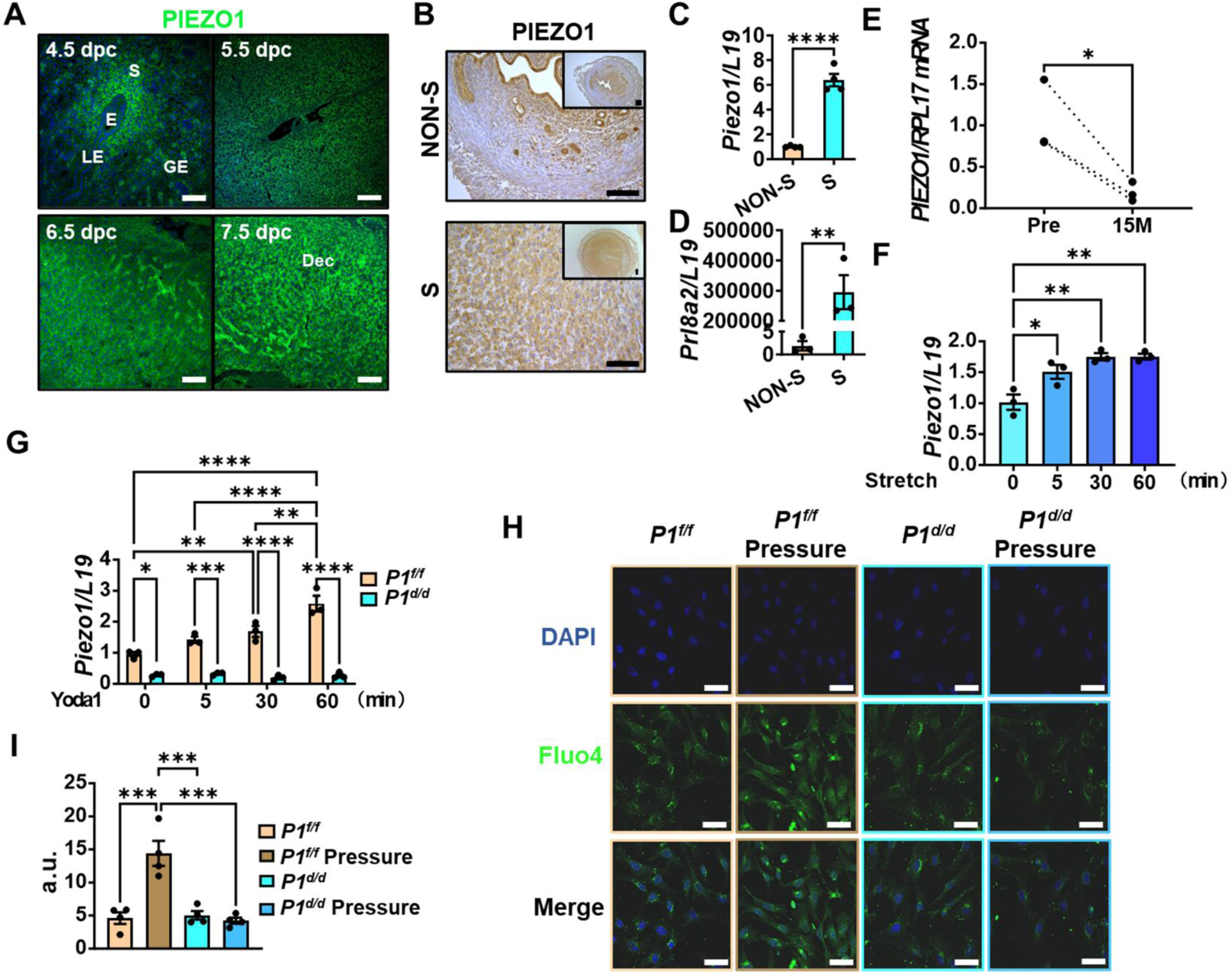
Expression of functional PIEZO1 in mouse and baboon endometrium. (A) Expression of PIEZO1-tdTomato fusion protein in implantation sites at 4.5, 5.5, 6.5, and 7.5 dpc in *Piezo1^tdTomato^* mice detected by anti-RFP antibody. S, stromal cells; IS, implantation sites; GE, glandular epithelial cells; (B) Expression of PIEZO1 in non-stimulated (NON-S) and artificial decidualized (S) uterine horns of WT mice, detected by anti-PIEZO1 antibody; (C) qPCR detected the mRNA level of *Piezo1* in non-stimulated (NON-S) and artificial decidualized (S) uterine horns of WT mice; (D) qPCR detected mRNA level of decidualization marker *Prl8a2* in non-stimulated (NON-S) and artificial decidualized (S) uterine horns of WT mice; (E) Expression of *PIEZO1* in eutopic baboon endometrium before (Pre) and 15 months after induction of endometriosis. (F) qPCR detected the mRNA level of *Piezo1* in *ex vivo* cultured uterine tissue of WT mice after mechanical stretching (0, 5, 30, and 60 min); (G) qPCR detected the mRNA level of *Piezo1* in *ex vivo* cultured uterine tissue of *Piezo1^f/f^* (*P1^f/f^*) and *Piezo1^d/d^* (*P1^d/d^*) mice after treating with Yoda1 (0, 5, 30, and 60 min); (H) Intracellular Ca^2+^ levels of mESCs isolated from *Piezo1^f/f^* (*P1^f/f^*) and *Piezo1^d/d^* (*P1^d/d^*) mice with (Pressure) or without 25 kPa of hydrostatic pressure detected by Fluo-4 Ca^2+^ probe (Green); (I) The quantification of fluorescence in F; Scale bar = 100 μm in A&B; Scale bar = 50 μm in H; *p<0.05; **p<0.01; ***p<0.001; ****p<0.0001.

To assess the functional activity of PIEZO1 in medicating Ca^2+^ influx in uterine cells, we crossed *Piezo1^f/f^* mice with *Pgr^Cre/+^* mice to generate a uterine-specific *Piezo1* knockout mouse model (*Pgr^Cre/+^Piezo1^f/f^*, referred to as *Piezo1^d/d^*). The littermates without Pgr-Cre (*Pgr^+/+^Piezo1^f/f^*, referred to as *Piezo1^f/f^*) were used as a control (Fig. S1B). The absence of PIEZO1 in the endometrium was verified through qPCR and immunohistochemistry (Fig. S1C&D). Next, we applied a hydrostatic pressure of 25 kPa onto *in vitro* cultured mouse stromal cells (mESCs). We detected increased cytoplastic Ca^2+^ levels using the fluo-4 AM Ca^2+^ probe in mESCs isolated from *Piezo1^f/f^* mice but not in mESCs from the *Piezo1^d/d^* mice, confirming the functioning of PIEZO1 protein in these cells (Fig. 1H&I). Collectively, these findings confirm the expression of functional PIEZO1 in the mouse endometrium, specifically in stromal cells during decidualization.

### Piezo1 is critical for decidualization in the endometrium

Next, we investigated the impact of PIEZO1 on mouse pregnancy by mating the *Piezo1^f/f^* and *Piezo1^d/d^* females with fertile wild-type males. A four-month fertility test showed significantly smaller litter size and number of litters/mouse in the *Piezo1^d.d^* mice compared to *Piezo1^f/f^* mice (Figure 2A). Notably, the absence of PIEZO1 did not exert any influence on the number and weight of implantation sites at 7.5 dpc during early pregnancy (Fig. 2B-D). However, at 11.5 and 14.5 dpc, a considerable number of embryos were absorbed in the *Piezo1^d/d^*uterus, suggesting the presence of a developmental disorder during mid-late pregnancy (Fig. 2B-D). The weight of the remaining fetus, not the placentas, was significantly lower in the *Piezo1^d.d^* mice compared to *Piezo1^f/f^* mice at 14.5 dpc (Fig. S1E&F). We, therefore, conducted a comparison of the decidua between *Piezo1^f/f^* and *Piezo1^d/d^* mice at 11.5 and 14.5 dpc. The histological staining revealed a significantly thinner decidua basalis in the uterine tissue lacking PIEZO1 expression at both time points (Fig. 3A). Moreover, qPCR results demonstrated significantly reduced levels of decidualization markers *Prl8a2*, *Prl3c1*, *Bmp2*, and *Wnt4* in the decidua tissue from the *Piezo1^d/d^* mice, indicating an impaired decidualization status in these mice (Fig. 3B). More importantly, treatment of Yoda1 in wildtype mice significantly up-regulated the expression of decidualization markers at 11.5 dpc (Fig. 3C).

**Figure 2.**
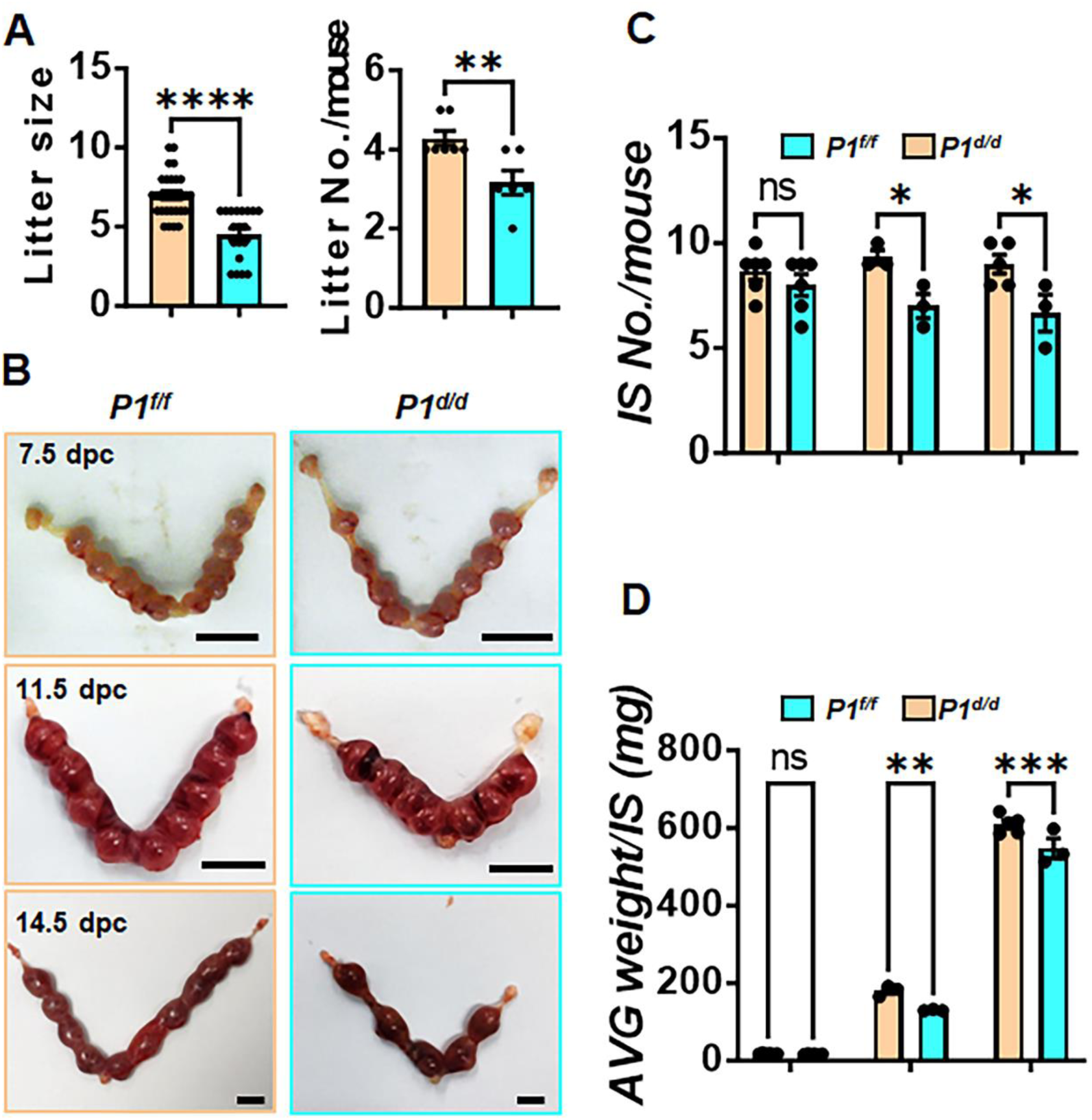
The pregnancy disorder of *Piezo1^d/d^* mice. (A) Litter size and number of litters per mouse of *Piezo1^f/f^* (*P1^f/f^*) and *Piezo1^d/d^* (*P1^d/d^*) mice in a 4-mouth fertility test; (B) Morphology of uterine horns of *Piezo1^f/f^* (*P1^f/f^*) and *Piezo1^d/d^* (*P1^d/d^*) mice at 7.5, 11.5 and 14.5 dpc; (C) Number of implantation sites of *Piezo1^f/f^* (*P1^f/f^*) and *Piezo1^d/d^* (*P1^d/d^*) mice at 7.5, 11.5 and 14.5 dpc; (D) Average weight of implantation sites from each mouse of *Piezo1^f/f^* (*P1^f/f^*) and *Piezo1^d/d^* (*P1^d/d^*) mice at 7.5, 11.5 and 14.5 dpc; Scale bar = 1 cm; ns p>0.05; *p<0.05; **p<0.01; ***p<0.001; ****p<0.0001.

**Figure 3.**
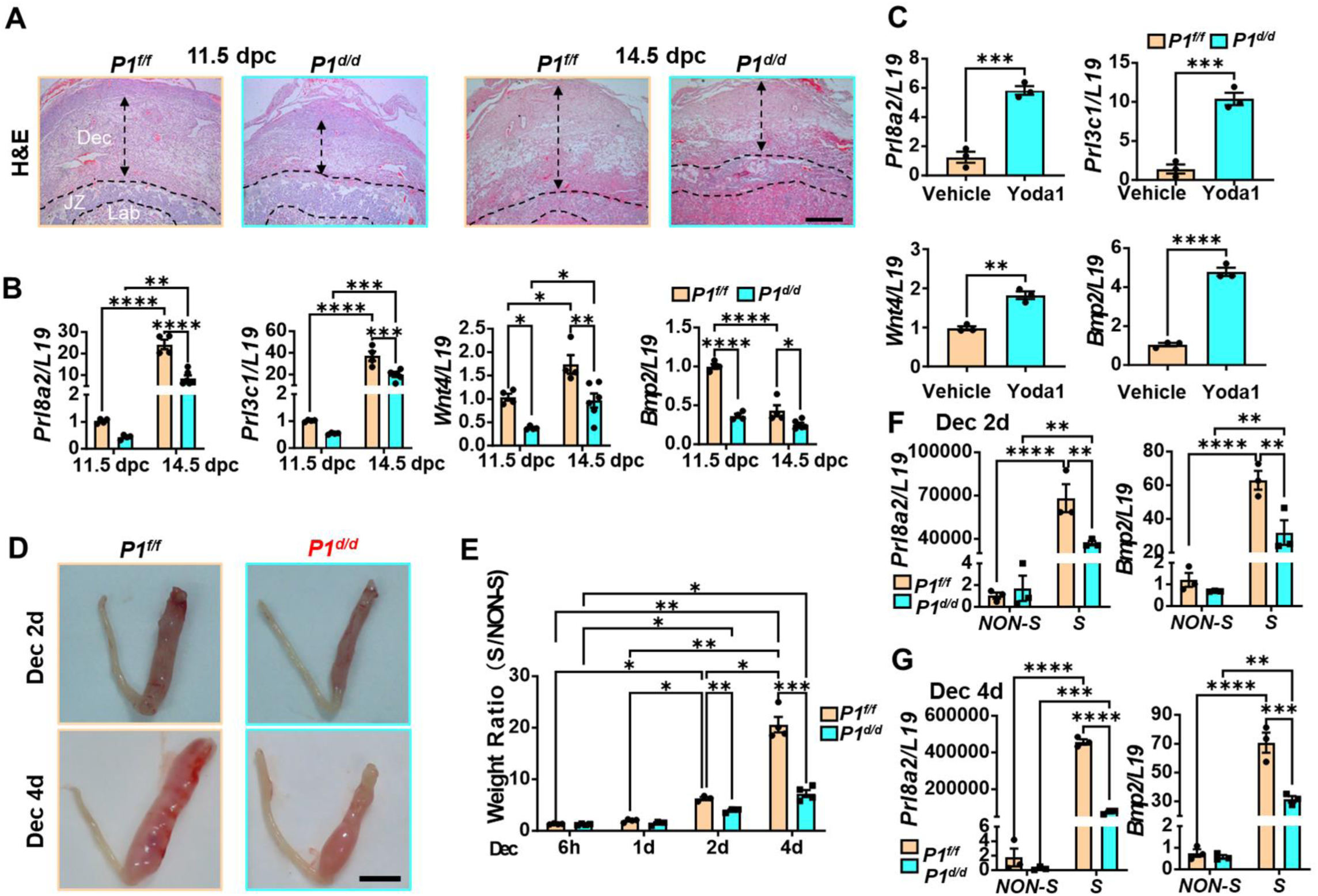
Critical roles of PIEZO1 to endometrial decidualization in mice and humans. (A) Histological images of decidua (Dec), junction zone (JZ), and labyrinth (Lab) of *Piezo1^f/f^* (*P1^f/f^*) and *Piezo1^d/d^* (*P1^d/d^*) mice at 11.5 and 14.5 dpc; (B) mRNA expression of decidualization marker genes in decidua of *Piezo1^f/f^* (*P1^f/f^*) and *Piezo1^d/d^* (*P1^d/d^*) mice at 11.5 and 14.5 dpc; (C) mRNA expression of decidualization marker genes in the decidua of WT mice with (Yoda1) or without (Vehicle) Yoda1 treatment; (D) Morphology of stimulated (S) and non-stimulated (NON-S) uterine horns of artificially decidualized *Piezo1^f/f^* (*P1^f/f^*) and *Piezo1^d/d^* (*P1^d/d^*) mice at 2 and 4 days after stimulation (E) Ratio of the uterine wet weight of artificially decidualized *Piezo1^f/f^* (*P1^f/f^*) and *Piezo1^d/d^* (*P1^d/d^*) mice 6 hours, 1, 2, and 4 days after stimulation; (F) mRNA expression of decidualization marker genes in deciduma of *Piezo1^f/f^* (*P1^f/f^*) and *Piezo1^d/d^* (*P1^d/d^*) mice 2 days after stimulation; (G) mRNA expression of decidualization marker genes in deciduma of *Piezo1^f/f^* (*P1^f/f^*) and *Piezo1^d/d^* (*P1^d/d^*) mice 4 days after stimulation; Scale bar = 500 μm in A; Scale bar = 1 cm in D; *p<0.05; **p<0.01; ***p<0.001; ****p<0.0001.

To further validate the crucial role of PIEZO1 in regulating decidualization, while isolating the influence of embryo and ovaries, we employed an artificial decidualization model that induces decidualization through a single mechanical stimulation in ovariectomized mice primed with exogenous E_2_ and P_4_ [40]. One day after mechanical stimulation, the levels of *Prl8a2* were notably lower in *Piezo1^d/d^* mice compared to *Piezo1^f/f^* mice, despite no differences observed in the weight of deciduoma (Fig. 3D, Fig. S2G). Subsequently, at 2 and 4 days post-stimulation, the weight of the stimulated horn in the *Piezo1^f/f^* mice increased more than 6.2 and 20.4 times, respectively, in comparison to the non-stimulated control horn (Fig. 3D&E). In contrast, the deciduoma showed only approximately 3.9 and 7.2 times greater weight compared to the control horns in the *Piezo1^d/d^* mice, respectively (Fig. 3D&E). Similarly, the expression levels of decidualization markers were significantly lower in the stimulated horns of the *Piezo1^d/d^* mice than that of the *Piezo1^f/f^* mice, indicating the impaired artificial decidualization in the absence of PIEZO1 (Fig. 3F&G). Collectively, these findings strongly suggest that the absence of PIEZO1 disrupts the development, but not the initiation of decidualization, consistent with our observation during the natural pregnancy.

Next, we investigated whether PIEZO1 plays a favorable role in the *in vitro* decidualization process of mESCs. When exposed to the well-established cocktail of E_2_+P_4_ to induce decidualization, mESCs derived from *Piezo1^d/d^* mice exhibited a notable reduction in the expression of decidualization markers *Prl8a2* and *Prl3c1* compared to mESCs from *Piezo1^f/f^* mice (Fig. 4A&B). Co-treatment with Yoda1 significantly enhanced the decidualization response in mESCs isolated from *Piezo1^f/f^* mice (Fig. 4A&B). This Yoda1-enhanced decidualization effect was abolished in mESCs isolated from *Piezo1^d/d^*mice (Fig. 4A&B). Furthermore, in human endometrial stromal cells (HESCs), silencing *PIEZO1* expression by siRNA resulted in a significantly decreased expression of decidualization markers *FOXO1*, *PRL*, and *IGFBP1* following treatment with the decidualization-inducing cocktail MPA+cAMP (Fig. 4C-E). Notably, co-treatment with Yoda1 significantly enhanced the expression of all three decidualization marker genes in siCtrl-transfected but no such effect was observed in siPIEZO1-transfected HESCs (Fig. 4C-E). These findings demonstrate the conserved role of PIEZO1 in supporting successful decidualization across humans and mice.

**Figure 4.**
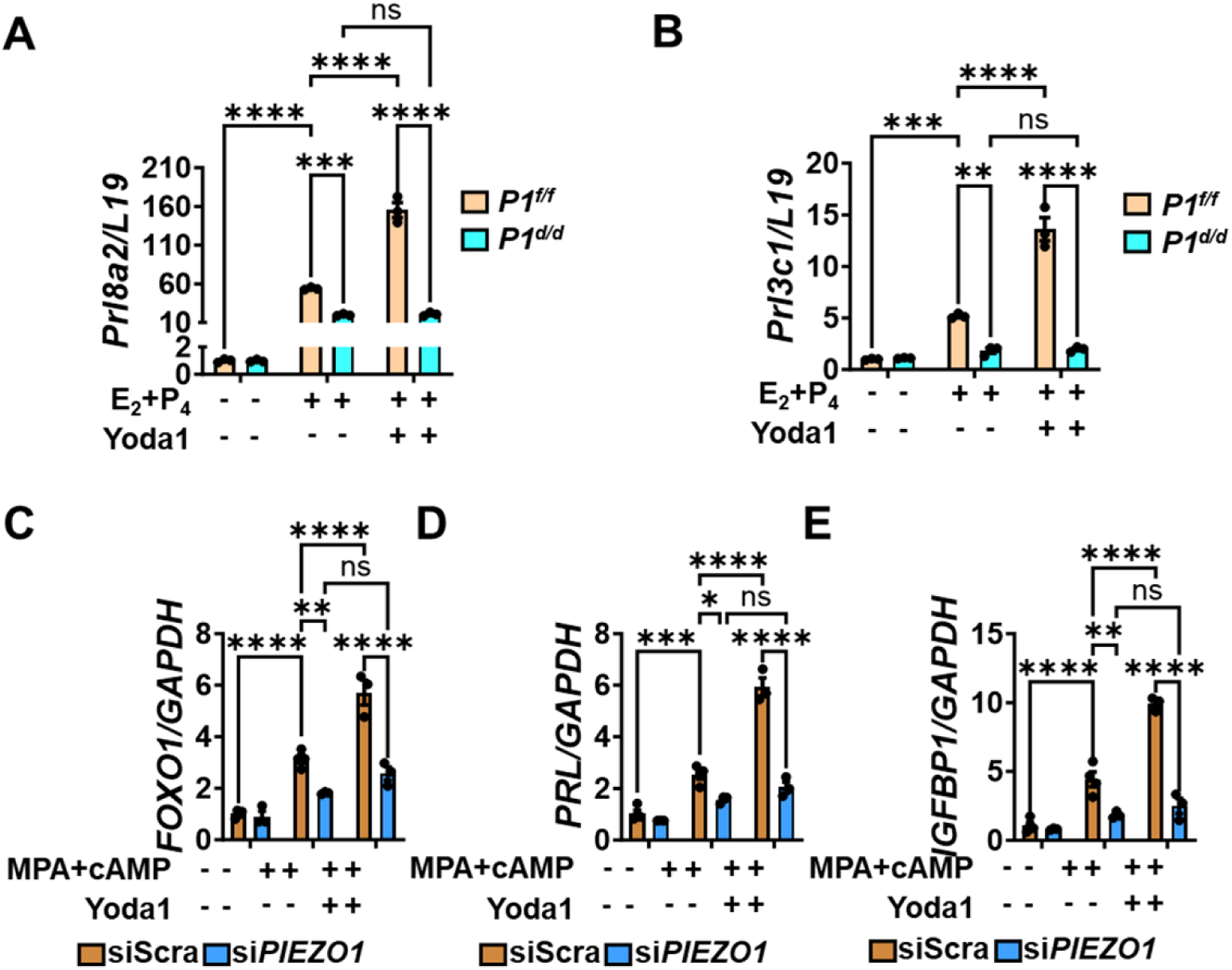
Critical roles of Yoda1/PIEZO1 to *in vitro* decidualization in mESCs and HESCs. (A&B) mRNA expression of decidualization marker genes in 2-day *in vitro* decidualized mESCs (E_2_+P_4_ treatment) isolated from *Piezo1^f/f^* (*P1^f/f^*) and *Piezo1^d/d^* (*P1^d/d^*) mice with or without co-treatment with Yoda1; (C-E) mRNA expression of decidualization marker genes in 2-day *in vitro* decidualized, scrambled (Ctrl) or *PIEZO1* siRNA (P1-KD) transfected HESCs (MPA+cAMP treatment), with or without co-treatment with Yoda1; ns p>0.05; *p<0.05; **p<0.01; ***p<0.001; ****p<0.0001.

### Stromal cell contraction and ECM stiffness affect decidualization via activation of PIEZO1

Previous studies have demonstrated that stromal cells exhibit a myofibroblastic phenotype, termed fibroblast activation, during the early stage of decidualization in human, baboon, and mouse models, which is characterized by the increased expression of the cytoskeletal protein α smooth muscle actin (αSMA), indicating enhanced contraction capability of these cells [41–43]. Consistent with this, our results revealed that decidualization resulted in significantly increased contractility, with the gel area reduced by about 50.8% in mESCs derived from *Piezo1^f/f^* mice, compared to a reduction of approximately 23.1% in non-decidualized cells (Fig. 4A&B). Interestingly, mESCs driven from *Piezo1^d/d^* mice exhibited a notably lower contraction during decidualization, with the gel area decreasing by 39.9%, accompanied with their impaired decidualization status (Fig. 5A&B). However, treatment with butanedione monoxime (BDM), an inhibitor of cell contraction, effectively suppressed the decidualization-induced contraction of *Piezo1^f/f^* mESCs but did not on *Piezo1^d/d^* mESCs (Fig. 5A&B). Moreover, BDM treatment significantly impaired the *in vitro* decidualization of cells isolated from *Piezo1^f/f^* mice, but not cells from *Piezo1^d/d^*mice (Fig. 5C). Collectively, these findings suggest that the enhanced contractility of stromal cells during decidualization activates PIEZO1, thereby promoting decidualization in mice, establishing a positive feedback loop.

**Figure 5.**
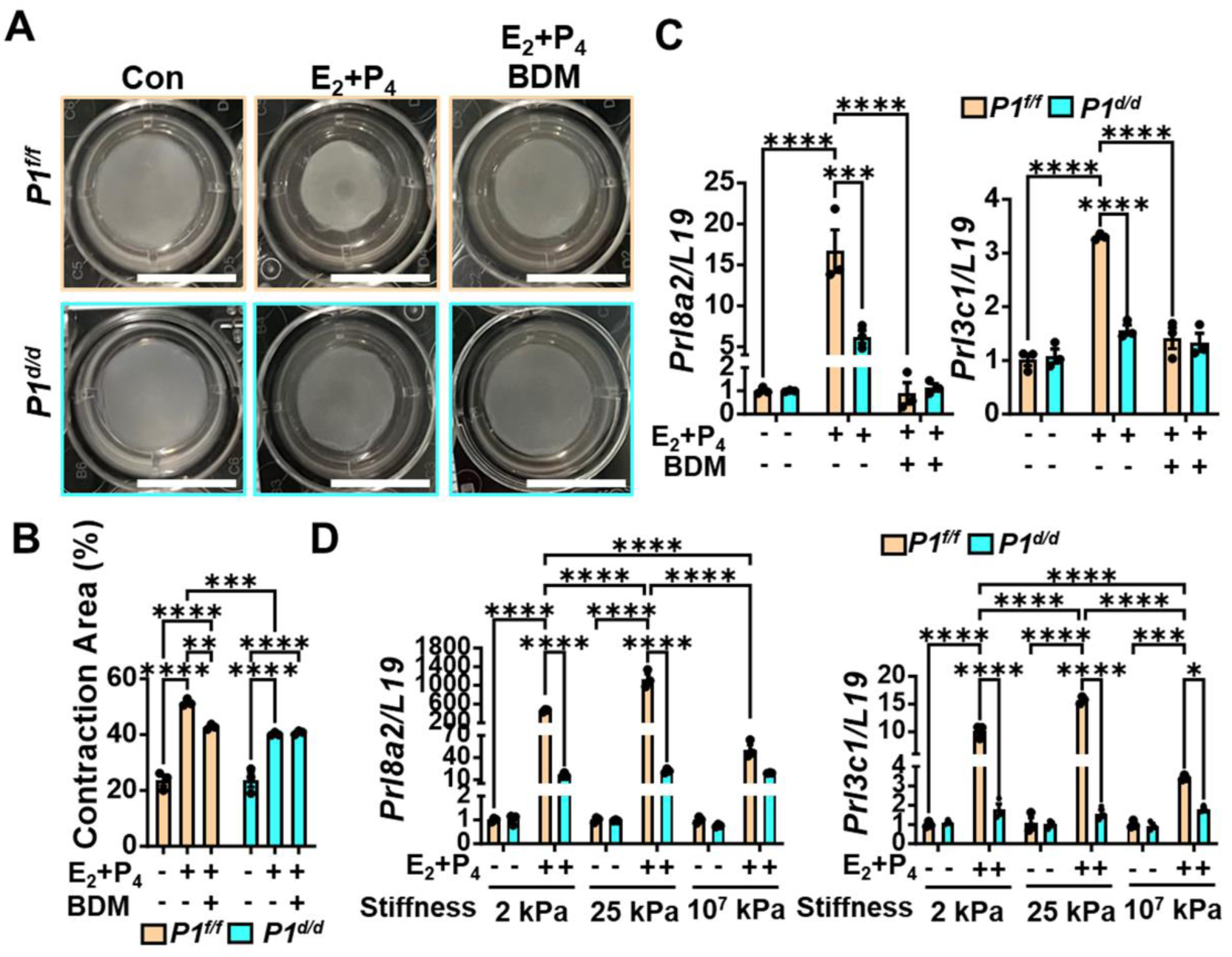
Cell contraction and ECM stiffness affect decidualization through PIEZO1. (A) Collagen contraction assay of *in vitro* decidualized mESCs (E_2_+P_4_ treatment, EP) isolated from *Piezo1^f/f^* (*P1^f/f^*) and *Piezo1^d/d^* (*P1^d/d^*) mice, the vehicle was used as control (CON), butanedione monoxime (BDM) was used as an inhibitor of cell contraction; (B) The quantification of contraction area (%) in A; (C) mRNA expression of decidualization marker genes in 2-day *in vitro* decidualized mESCs (E_2_+P_4_ treatment) isolated from *Piezo1^f/f^*(*P1^f/f^*) and *Piezo1^d/d^* (*P1^d/d^*) mice with or without co-treatment with BDM; (D) mRNA expression of decidualization marker genes in 2-day *in vitro* decidualized mESCs (E_2_+P_4_ treatment) isolated from *Piezo1^f/f^*(*P1^f/f^*) and *Piezo1^d/d^* (*P1^d/d^*) mice cultured on the surface with different stiffness. Scale bar = 1 cm; *p<0.05; **p<0.01; ***p<0.001; ****p<0.0001.

The elasticity or stiffness of ECM influences numerous fundamental cellular processes, including proliferation, differentiation, and organoid formation [44]. The endometrium undergoes ECM rapid remodeling during decidualization [8]. Therefore, we examined the impact of different ECM stiffness on the process of decidualization. We seeded the mESCs on Softwell^®^ plates with varying stiffness (2 kPa and 25 kPa), coated with type I collagen (COL I), while a regular petri dish, coated with COL I, served as high-stiffness control (∼ 10^7^ kPa). *Piezo1^f/f^* mESCs cultured on softer substrates (2 kPa and 25 kPa) exhibited considerably higher expression levels of decidualization markers (*Prl8a2* and *Prl3c1*) in response to *in vitro* decidualization, compared to those on high-stiffness control (Fig. 5D). In contrast, *Piezo1^d/d^* mESCs did not show any difference in decidualization markers across substrates with different stiffness, suggesting that ECM stiffness effectively regulates the decidualization response through PIEZO1 (Fig. 5D). These collective findings suggest that the contractility of stromal cells is activated during decidualization, interacts with ECM stiffness to activate PIEZO1, and eventually enhances decidualization.

### PIEZO1 regulates decidualization through the Ca^2+^-CaMKII cascade

Next, we investigated whether PIEZO1 influences stromal cell decidualization through its primary role as a Ca^2+^ ion channel modulator. The Fluo-4 AM fluorescence analysis showed a significant increase in cytoplastic Ca^2+^ levels in *Piezo1^f/f^* mESCs under *in vitro* decidualization conditions, whereas *Piezo1^d/d^*mESCs exhibited no such response, indicating that decidualization activates PIEZO1-mediated Ca^2+^ influx (Fig. 6A&B). Furthermore, upon treatment with Yoda1, a robust higher cytoplastic Ca^2+^ level was observed in the *Piezo1^f/f^* cells during *in vitro* decidualization, while *Piezo1^d/d^* cells failed to respond (Fig. 6A&B). To explore the relationship between Yoda1-induced Ca^2+^ level and the previously mentioned Yoda1-enhanced decidualization, we employed BAPTA-AM, a cell-permeating Ca^2+^ chelator, to block the intracellular Ca^2+^. The results demonstrated that BAPTA-AM significantly abolished Yoda1-enhanced decidualization in *Piezo1^f/f^* mESCs (Fig. 6C). Similarly, BAPTA-AM played the same role in HESCs (Fig. 6D). These findings indicate that PIEZO1 regulates decidualization by mediating Ca^2+^ influx in both mESCs and HESCs.

**Figure 6.**
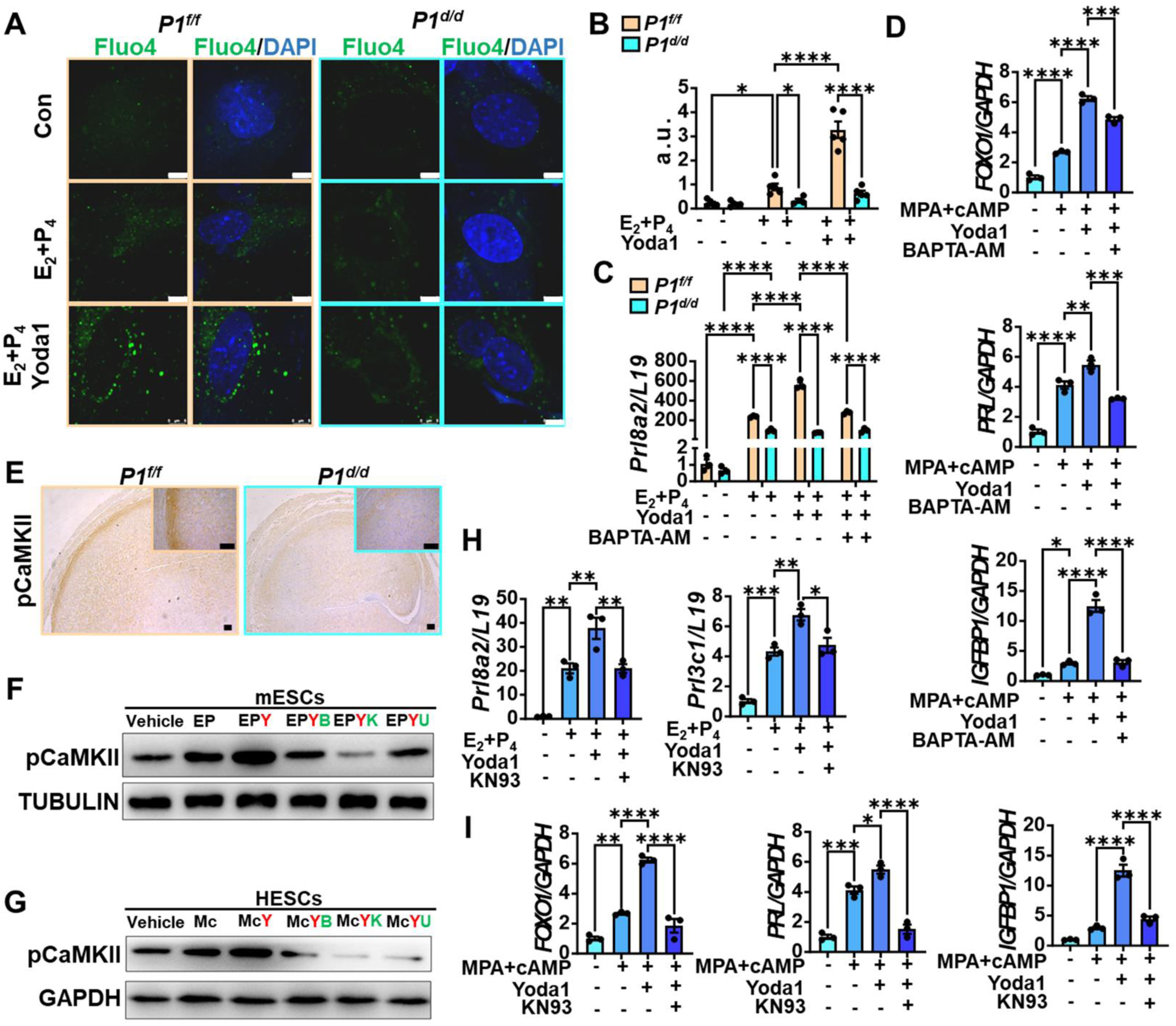
PIEZO1 enhances decidualization through the Ca^2+^-CamKII cascade. (A) Intracellular Ca^2+^ levels of in 2-day *in vitro* decidualized mESCs isolated from *Piezo1^f/f^* (*P1^f/f^*) and *Piezo1^d/d^*(*P1^d/d^*) mice co-treated with Yoda1, detected by Fluo-4 Ca^2+^ probe (Green); (B) The quantification of fluorescence in A; (C) mRNA expression of decidualization marker *Prl8a2* in 4-day *in vitro* decidualized mESCs (E_2_+P_4_ treatment) isolated from *Piezo1^f/f^* (*P1^f/f^*) and *Piezo1^d/d^* (*P1^d/d^*) mice, co-cultured with Yoda1 and BAPTA-AM; (D) mRNA expression of decidualization marker genes in 2-day *in vitro* decidualized HESCs (MPA+cAMP treatment), co-cultured with Yoda1 and BAPTA-AM; (E) Immunostaining of pCaMKII in artificial decidualized uterine horns of *Piezo1^f/f^* (*P1^f/f^*) and *Piezo1^d/d^* (*P1^d/d^*) mice 4 days after stimulation; (F) Expression of pCaMKII in 2-day *in vitro* decidualized mESCs (E_2_+P_4_ treatment, EP) co-treated with Yoda1 (Y), BAPTA-AM (B), KN93 (K), or U0126 (U), the vehicle was used as control; (G) Expression of pCaMKII in 2-day *in vitro* decidualized HESCs (MPA+cAMP treatment, Mc) co-treated with Yoda1 (Y), BAPTA-AM (B), KN93 (K), or U0126 (U); (H) mRNA expression of decidualization marker genes in 2-day *in vitro* decidualized mESCs (E_2_+P_4_ treatment), co-cultured with Yoda1 and KN93; (I) mRNA expression of decidualization marker genes in 2-day *in vitro* decidualized HESCs (MPA+cAMP treatment), co-cultured with Yoda1 and KN93; Scale bar = 5 μm in A; Scale bar = 100 μm in E; *p<0.05; **p<0.01; ***p<0.001; ****p<0.0001.

Calcium/calmodulin-dependent kinase II (CaMKII) functions as a downstream target of the Ca^2+^-calmodulin complex and plays a pivotal role in regulating many cellular functions [45], which retains its ability to undergo Ca^2+^-and calmodulin-dependent auto-phosphorylation at Thr286 or Thr287 [46, 47]. Inhibiting CaMKII activation down-regulates the expression of decidualization marker genes *PRL* and *IGFBP1* in HESCs [48]. Therefore, we hypothesized that CaMKII acts as a downstream mediator of Ca^2+^ influx induced by PIEZO1. Firstly, we evaluated the phosphorylation level of CaMKII in *in vivo* artificial decidualization model. The results revealed the increased expression of phosphorylated CaMKII (pCaMKII) in the secondary decidual zone (SDZ) of mechanically stimulated uterine horns of *Piezo1^f/f^*mice 4 days after stimulation but was significantly lower in *Piezo1^d/d^*mice (Fig. 6E). Moreover, compared to the vehicle-treated control group, the pCaMKII level was significantly up-regulated in both *Piezo1^f/f^* mESCs and HESCs under *in vitro* decidualization conditions and further enhanced by co-treatment with Yoda1 (Fig. 6F&G). In addition, when intracellular Ca^2+^ was chelated by BAPTA-AM, the Yoda1-induced pCaMKII phosphorylation was inhibited (Fig. 6F&G). These findings strongly suggest a close association between pCaMKII levels and Yoda1/PIEZO1 enhanced decidualization. To provide further evidence for the necessity of elevated pCaMKII in Yoda1-enhanced decidualization, we treated mESCs and HESCs with KN93, a pCaMKII inhibitor, in combination with Yoda1 and the decidualization-inducing cocktail. Remarkably, the expression of decidualization markers induced by Yoda1 treatment was significantly reversed by KN93, indicating that pCaMKII mediated Yoda1-enhanced decidualization (Fig. 6H&I). Taken together, our data provide evidence that Yoda1/PIEZO1 governs decidualization through the Ca^2+^-CaMKII cascade.

### PIEZO1 promotes decidualization via ERK1/2 MAPK signaling

Next, we employed RNA-Seq to analyze decidualized and non-decidualized uterine horns from both *Piezo1^f/f^* and *Piezo1^d/d^* mice 4 days after artificial decidualization. Differentially expressed genes (DEGs) meeting the criteria of a fold change greater than 2.0 and an adjusted p-value lower than 0.05 were selected from four distinct comparisons as depicted in Fig. S2A. In non-decidualized uterine horns, *Piezo1* knockout resulted in only 78 significantly up-regulated genes and 47 down-regulated genes (Comparison 1, Fig. S2B), whereas decidualized horns exhibited significant changes, with 1364 up-regulated and 551 down-regulated genes (Comparison 2, Fig. S2C). These data suggest that the absence of *Piezo1* resulted in minimal changes in non-decidualized uterine horns but significant alterations in gene expression during the decidualization. Further analysis compared the differences of decidualization on gene expression in *Piezo1^f/f^*and *Piezo1^d/d^* mice, which were shown in comparison 3 and 4, respectively, provided additional insights. (Fig. S2D&E). By overlapping of DEGs of comparisons 2 and 3 visualized in a Venn diagram (Fig. S2F), we identified 1474 genes that were significantly altered by both decidualization and the absence of *Piezo1* (Fig. S2G). KEGG pathway analysis of these 1474 DEGs highlighted multiple affected signaling pathways, including the calcium signaling pathway, which aligns with the primary function of PIEZO1 as previously described (Fig. 7A). Enriched pathways also included cAMP, Wnt, Rap1, and its downstream MAPK signaling pathways (Fig. 7A), all known to be critical for successful decidualization [48, 49].

**Figure 7.**
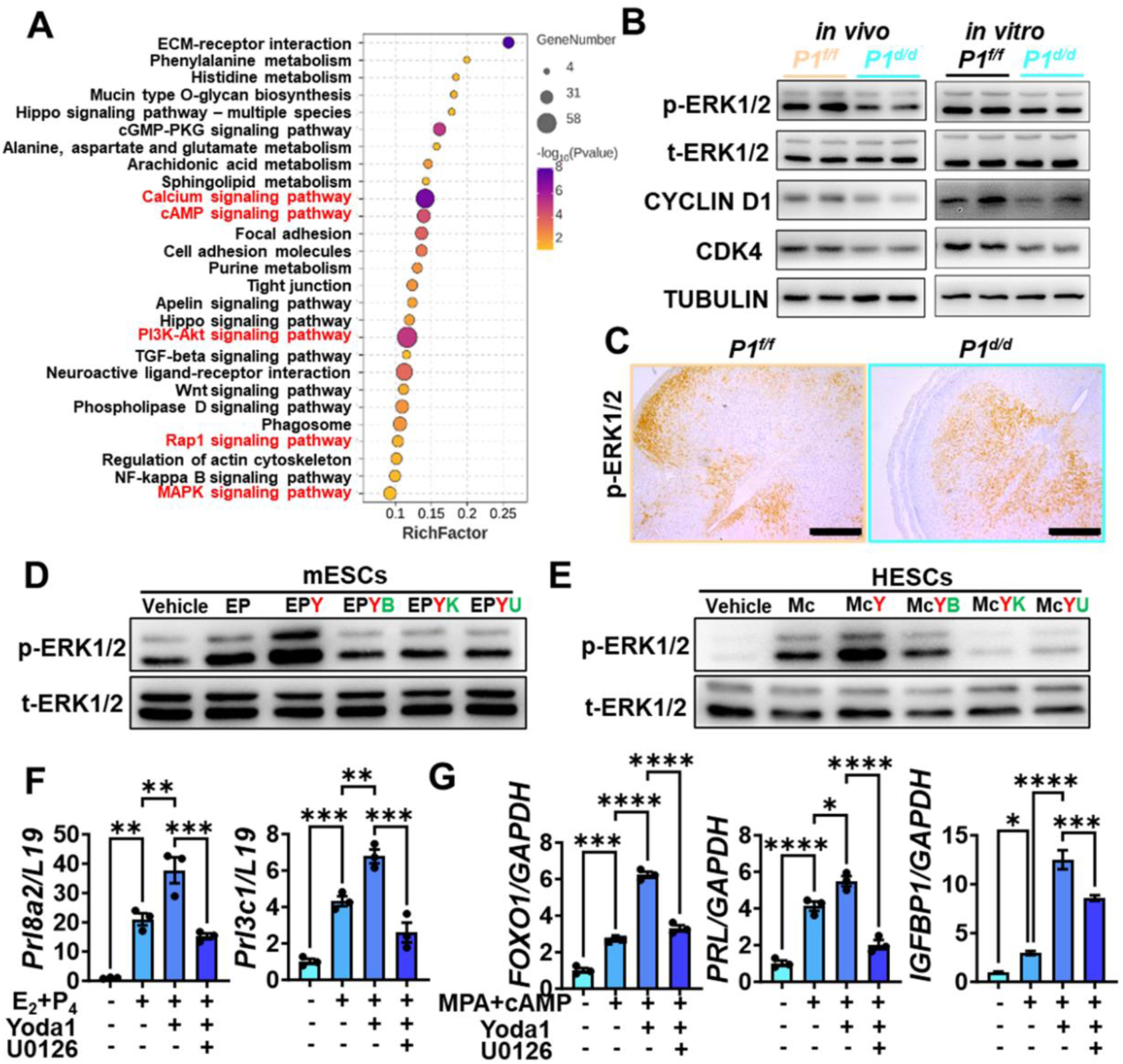
PIEZO1 enhances decidualization through MAPK signaling. (A) KEGG pathway analysis of DEGs between artificial decidualized uteri from *Piezo1^f/f^* (*P1^f/f^*) and *Piezo1^d/d^*(*P1^d/d^*) mice 4 days after stimulation; (B) Expression of phosphorylated ERK1/2 (p-ERK1/2), total ERK1/2 (t-ERK1/2), CYCLIN D1, and CDK4 in artificial decidualized uteri (*in vivo*, 4 days) and *in vitro* decidualized mESCs (*in vitro*, 2 days) of *Piezo1^f/f^* (*P1^f/f^*) and *Piezo1^d/d^* (*P1^d/d^*) mice; (C) Immunostaining of p-ERK1/2 in artificial decidualized uterine horns of *Piezo1^f/f^* (*P1^f/f^*) and *Piezo1^d/d^* (*P1^d/d^*) mice 4 days after stimulation; (D) Expression of p-ERK1/2 in 2-day *in vitro* decidualized mESCs (E_2_+P_4_ treatment, EP) co-treated with Yoda1 (Y), BAPTA-AM (B), KN93 (K), or U0126 (U); (E) Expression of pERK1/2 in 2-day *in vitro* decidualized HESCs (MPA+cAMP treatment, Mc) co-treated with Yoda1 (Y), BAPTA-AM (B), KN93 (K), or U0126 (U); (F) mRNA expression of decidualization marker genes in 2-day *in vitro* decidualized mESCs (E_2_+P_4_ treatment), co-cultured with Yoda1 and U0126; (G) mRNA expression of decidualization marker genes in 2-day *in vitro* decidualized HESCs (MPA+cAMP treatment), co-cultured with Yoda1 and U0126; Scale bar = 500 μm; *p<0.05; **p<0.01; ***p<0.001; ****p<0.0001.

Previous study shows that the ERK1/2 MAPK pathway is required for endometrial decidualization[50]. Several studies have shown that calcium signaling induces cell differentiation through CaMKII and MAPK/ERK pathways [51–53]. Therefore, we further investigated the potential involvement of the ERK1/2 MAPK signaling pathway as a mediator of decidualization failure resulting from the absence of *Piezo1*. First, we examined the change in phosphorylated ERK1/2 levels between *Piezo1^f/f^* and *Piezo1^d/d^* mice during decidualization. Western blot confirmed a significant decrease in pERK1/2 levels in both *in vivo* and *in vitro* decidualized tissue/cells of *Piezo1^d/d^* mice compared to those of *Piezo1^f/f^* mice, which correlated with the reduced expression of cell cycle regulators CyclinD1 and CDK4 (Fig. 7B). Immunostaining further revealed distinct localization patterns of pERK1/2 between *Piezo1^f/f^* and *Piezo1^d/d^* mice. In *Piezo1^f/f^* decidualized horns, pERK1/2 showed higher expression in the SDZ of *Piezo1^f/f^* mice, whereas pERK1/2 was predominantly localized in the primary decidual zone (PDZ) in *Piezo1^d/d^* decidualized horns at 4 days post-stimulation (Fig. 7C). Additionally, we investigated whether Yoda1 enhanced decidualization via pERK1/2 activation. Results from the western blot and immunostaining demonstrated a significant induction of pERK1/2 levels and nuclear localization upon decidualization, which was further enhanced by Yoda1 treatment in decidualized stromal cells (Fig. 7D, S2H). Similarly, the pERK1/2 level was also induced by decidualization and further enhanced by Yoda1 in HESCs (Fig. 7E). Conversely, the induction was attenuated by MEK/ERK inhibitor U0126, accompanied by the decreased expression of decidualization markers in both mESCs and HESCs, indicating the necessity of pERK1/2 activation for Yoda1/PIEZO1-induced decidualization (Fig. 7F&G). Furthermore, BAPTA-AM or KN93 treatment significantly decreased pERK1/2 levels, indicating that ERK/MAPK signaling acts downstream of the Ca^2+^-CaMKII cascade during Yoda1/PIEZO1-enhanced decidualization (Fig. 7D&E). Interestingly, the Yoda1-induced pCaMKII level decreased by U0126 in HESCs but not mESCs, suggesting a feedback loop between Ca^2+^/CamKII and ERK/MAPK signaling in humans (Fig. 7F&G). All these data collectively prove that the activation of PIEZO1 enhances decidualization through the Ca^2+^-CaMKII-ERK pathway.

### Autophagy mediates Yoda1/PIEZO1 enhanced decidualization depending on pBECN1

Multiple studies have shown that the intracellular Ca^2+^, pCaMKII, and ERK1/2 MAPK signaling regulate autophagy by regulating the phosphorylation of BECN1, a core protein of the class III PI3K nucleation complex [54–58]. In our KEGG pathway analysis, the autophagy pathway is one of the top 15 significantly changed cell processes (Fig. S3A). Therefore, we hypothesized that autophagy might mediate Yoda1/PIEZO1 enhanced decidualization. We first analyzed the expression of autophagy-related proteins in our *in vivo* and *in vitro* models. The results showed that the level of LC3A/B, the protein that contributes to the formation of autophagosome, significantly decreased in the decidualized tissue of *Piezo1^d/d^* mice compared to that of *Piezo1^f/f^* mice (Fig. 8A). Additionally, transmission electron microscopy (TEM) showed that the number of autophagic vacuoles was markedly increased in *in vitro* decidualized *Piezo1^f/f^* mESCs compared to non-decidualized cells, but not *Piezo1^d/d^*cells (Fig. 8B&C). In addition, LC3-GFP-RFP adenoviral transfection was employed to measure autophagic flux. We observed significantly fewer autophagosomes (yellow dots) and autolysosomes (free red dots) in decidualized stromal cells from *Piezo1^d/d^* mice than that of *Piezo1^f/f^* mice (Fig. 8D, Fig. S3B). The protein level of LC3A/B was increased by *in vitro* decidualization and further enhanced by the Yoda1 treatment in both mESCs and HESCs (Fig. 8E&F). These data indicate that autophagy is enhanced in the stromal cells during decidualization, mediated by PIEZO1.

**Figure 8.**
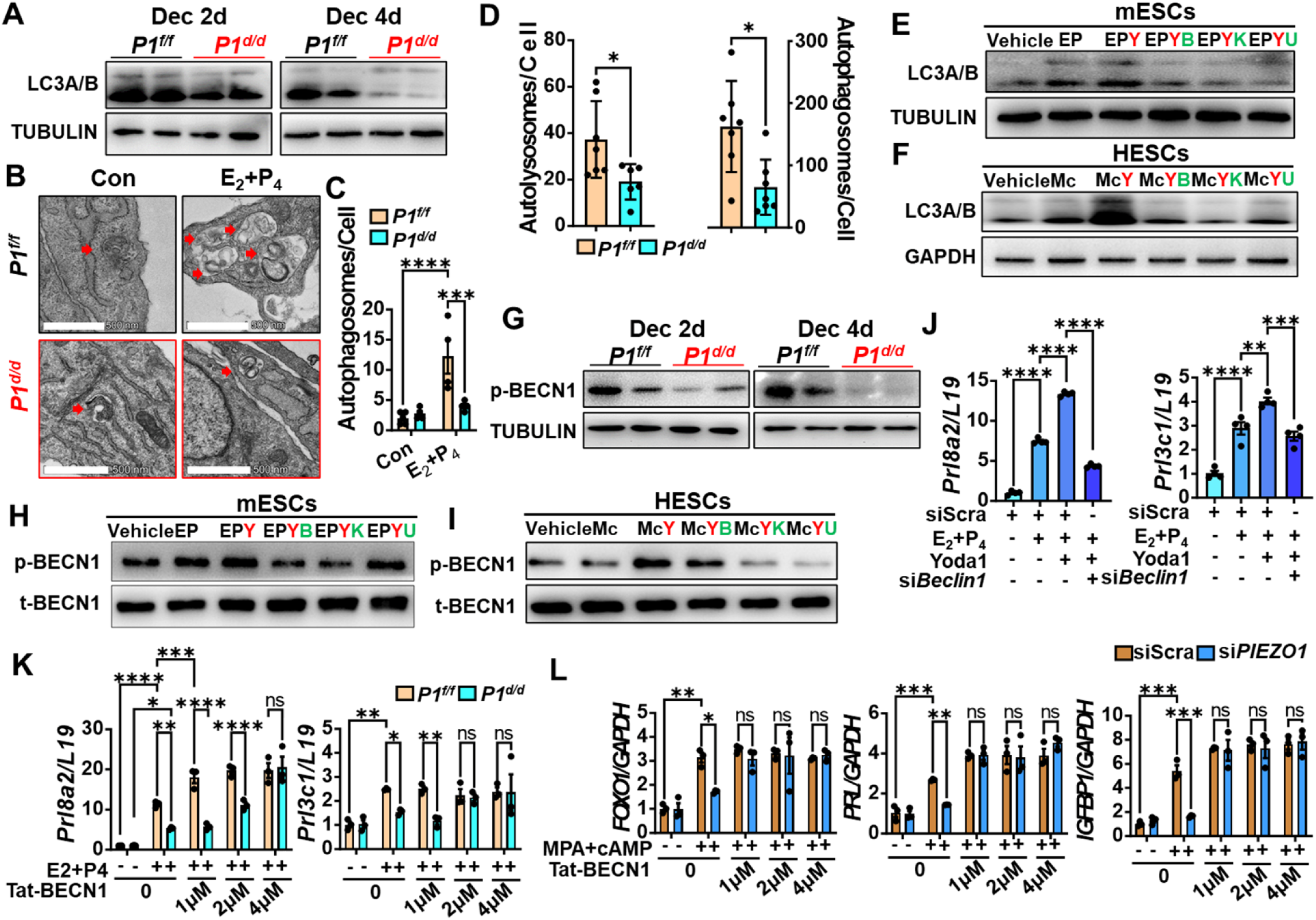
BECLIN1-dependent autophagy mediates Yoda1/PIEZO1 enhanced decidualization. (A) Expression of LC3A/B in artificial decidualized uteri of *Piezo1^f/f^* (*P1^f/f^*) and *Piezo1^d/d^* (*P1^d/d^*) mice 2 or 4 days after stimulation; (B) Representative TEM images of 2-day *in vitro* decidualized mESCs (E_2_+P_4_ treatment, EP) isolated from *Piezo1^f/f^* (*P1^f/f^*) and *Piezo1^d/d^* (*P1^d/d^*) mice, vehicle was used as control (CON), red arrows represent autophagosomes; (C) The quantification of autophagosomes in B; (D) The number of autophagosomes and autolysosomes in 2-day *in vitro* decidualized mESCs (E_2_+P_4_ treatment, EP) isolated from *Piezo1^f/f^*(*P1^f/f^*) and *Piezo1^d/d^* (*P1^d/d^*) mice, detected by LC3-GFP-RFP vector, representative images shown in figure S4A; (E) Expression of LC3A/B in 2-day *in vitro* decidualized mESCs (E_2_+P_4_ treatment, EP) co-treated with Yoda1 (Y), BAPTA-AM (B), KN93 (K), or U0126 (U); (F) Expression of LC3A/B in 2-day *in vitro* decidualized HESCs (MPA+cAMP treatment, Mc) co-treated with Yoda1 (Y), BAPTA-AM (B), KN93 (K), or U0126 (U); (G) Expression of p-BECN1 in artificial decidualized uteri of *Piezo1^f/f^* (*P1^f/f^*) and *Piezo1^d/d^* (*P1^d/d^*) mice 2 or 4 days after stimulation; (H) Expression of p-BECN1 in 2-day *in vitro* decidualized mESCs (E_2_+P_4_ treatment, EP) co-treated with Yoda1 (Y), BAPTA-AM (B), KN93 (K), or U0126 (U); (I) Expression of p-BECN1 in 2-day *in vitro* decidualized HESCs (MPA+cAMP treatment, Mc) co-treated with Yoda1 (Y), BAPTA-AM (B), KN93 (K), or U0126 (U); (J) mRNA expression of decidualization marker *Prl8a2* in 2-day *in vitro* decidualized, scrambled (siScra) or *Beclin1* siRNA (siBeclin1) transfected mESCs (E_2_+P_4_ treatment) isolated from *Piezo1^f/f^* (*P1^f/f^*) and *Piezo1^d/d^* (*P1^d/d^*) mice, co-cultured with or without Yoda1; (K) mRNA expression of decidualization marker *Prl8a2* in 2-day *in vitro* decidualized mESCs (E_2_+P_4_ treatment) isolated from *Piezo1^f/f^* (*P1^f/f^*) and *Piezo1^d/d^* (*P1^d/d^*) mice, co-cultured with different dose of Tat-BECLIN1; (L) mRNA expression of decidualization marker genes in 2-day *in vitro* decidualized HESCs (MPA+cAMP treatment), co-cultured with different doses of Tat-BECLIN1; Scale bar = 500 nm; ns p>0.05; *p<0.05; **p<0.01; ***p<0.001; ****p<0.0001.

Next, we further explored if Yoda1/PIEZO1 enhances decidualization by regulating autophagy through Ca^2+^/pCaMKII and pERK1/2 controlled BECN1-dependent manners. BECN1 has been shown to be phosphorylated at its Ser90 site by pCaMKII and subsequently promotes autophagy and cell differentiation in Hela cells [55]. Western blot analysis revealed that the level of p-BECN1 (Ser90) was significantly decreased in *Piezo1^d/d^* mice compared to the *Piezo1^f/f^* mice, following artificial decidualization (Fig. 8G). On the other hand, the Yoda1 treatment remarkably enhanced the p-BECN1 level in decidualized mESCs and HESCs, and was abolished by co-treatment with BAPTA-AM or KN93, suggesting Yoda1-induced p-BECN1 via Ca^2+^-pCaMKII cascade in both mouse and humans (Fig. 8H&I). However, the MEK/ERK inhibitor U0126 neutralized the Yoda1-enhanced LC3A/B levels, but was not able to affect Yoda1-induced p-BECN1 levels in mESCs, suggesting ERK/MAPK pathway-regulated autophagy in a BECIN1 Ser90 phosphorylation independent manner in mice (Fig. 8H). In contrast, U0126 inhibited both LC3A/B and pBECN1 in HESCs, consistent with its inhibition in pCaMKII level in these cells, suggesting difference in roles of ERK/MAPK signaling in regulating autophagy between humans and mice (Fig. 8I). When *Becn1* is silenced by siRNA, the decidualization-induced and Yoda1-enhanced autophagy was completely inhibited, evidenced by the expression level of LC3A/B (Fig. S3D). Most importantly, the Yoda1-enhanced decidualization marker expression was abolished by the silencing of *Becn1* (Fig. 8J). Conversely, the autophagy activator Tat-BECN1, a cell membrane permeable peptide derived from a region of BECN1 protein [59], was able to rescue the impaired decidualization in *Piezo1^d/d^* mESCs and si*PIEZO1-*treated HESCs (Fig. 8K&L). Interestingly, the level of p-mTOR was significantly decreased in decidualized *Piezo1^d/d^* stromal cells compared to that of *Piezo1^f/f^*cells under the same condition, suggesting an enhanced ULK1 complex activation in the absence of PIEZO1, which we considered as feedback due to inhibited autophagy process in these mice (Fig. S3C). Nevertheless, these data collectively indicates that autophagy mediates the Yoda1/PIEZO1 enhanced decidualization through the Ca^2+^-CaMKII-pBECN1 cascade in both mouse and human endometrium.

## Discussion

The decidua, is a key compartment of endometrium in humans, non-human primates, and rodents, which plays a critical role in various processes associated with pregnancy, including facilitating embryo implantation, selecting healthy embryos, and controlling trophoblast invasion[1]. Herein, we report PIEZO1 mediates mechanical force as a novel regulator of decidualization, in response to its microenvironment created by the contraction of stromal cells and the stiffness of ECM. Conditional knockout of *Piezo1* in the mouse uterus leads to impairment of decidualization in both natural pregnancy and artificial decidualization models. On the other hand, the PIEZO1 agonist Yoda1 also enhanced decidualization during pregnancy. *In vitro* cell culture experiments demonstrates that the role of PIEZO1 in decidualization is similar between humans and mice. Mechanistically, upon PIEZO1 channel activation by cell contraction and ECM stiffness, intracellular Ca^2+^ influx is generated which subsequently phosphorylates CaMKII. Further investigation demonstrated that PIEZO1-mediated Ca^2+^-CaMKII regulates decidualization through BECN1-dependent autophagy.

In this study, we first assessed the upstream mechanical cues that induce PIEZO1 activation. Cells exert intrinsic forces on their environment, that is, on the ECM and neighboring cells, through mechanisms such as actomyosin contractility and cytoskeletal assembly [60]. On the other hand, cell contraction can induce long-range stress stiffening in the ECM [61]. Multiple studies reveal that such mechanical forces direct stem cell behaviors in development and regeneration (reviewed in [60]. During decidualization, the endometrial stromal cells undergo differentiation, associated with rapid remodeling of the cytoskeleton and the surrounding ECM [8, 43]. Our data also shows that the contractility of decidualized stromal cells is stronger than non-decidualized ones. In humans, the stiffness of decidua basalis is significantly higher than that of non-decidualized endometrium [24]. These observations indicate that the decidualization of stromal cells can largely change their mechanical microenvironment, which in turn enhances decidualization via the activation of PIEZO1 proteins. Such inference may explain why the null phenotype of PIEZO1 impairs decidualization at a late stage, but not during the initiation of the process. Similarly, the decidua basalis at 11.5 dpc and 14.5 dpc are impaired by the absence of PIEZO1, but 7.5 dpc is not significantly affected. In addition, the invasion of trophoblastic cells can be another contributor to the changes of tissue stiffness in the decidua basalis [24]. Therefore, the fact that expression of *Piezo1* is up-regulated by stretch and Yoda1 treatment suggests positive feedback on PIEZO1 and its function during this process.

The endometrium undergoes ECM remodeling during decidualization, suggesting ECM may play an important role in the process [8]. The average stiffness of the human decidua basalis is much higher than endometrium from non-pregnant women [24]. Our study reports that mESCs growing on ECM of 25 kPa stiffness display a better decidualization response compare to the hard surface of regulator petri dishes, which confirms the importance of ECM stiffness in regulating decidualization. Similarly, a recent study showed significantly increased PRL secretion from decidualizing HESCs cultured in 3-D in soft PEG gels compared to cells cultured in 2D on the hard surface of regulator petri dishes [62]. More importantly, we demonstrate that PIEZO1 is the key modulator of ECM stiffness regulation on decidualization response. On the other hand, the mESCs growing on ECM with 2 kPa stiffness display a weakened decidualization response than mESCs on 25 kPa surface, which supports the finding that PIEZO channels are more prone to activation on rigid substrates with stiffness ranging from 11 to 30 kPa [22]. However, the average stiffness of the human decidua basalis is 1.25 kPa, and in some cases up to 6 kPa, which is lower than the PIEZO channel activation range and the stiffness used in our study [24], which may be due to the limitation of AFM measurements in the study, which measures the stiffness of tissue rather than the ECM. However, the fact that cells grown on 2 kPa and hard surface regulator petri dishes also display impaired decidualization when knocking out *Piezo1*, suggests that this channel can be activated within a large range of stiffness. Nevertheless, our data show that a certain stiffness is beneficial for decidualization through activation of PIEZO1. In addition, an *in vitro* study using a microfluidic model reported that hemodynamic forces activate PIEZO1 in endothelial cells, inducing endothelial-derived prostaglandin E2 and prostacyclin which subsequently enhances decidualization via a paracrine response, providing a different cascade through which the PIEZO1 channel contributes to decidualization [63].

It is well established that the PIEZO1 mechanotransduction mechanism permits a calcium influx that further modulates downstream intracellular pathways [64]. Indeed, using a Fluo-4 AM Ca^2+^ probe, we showed that PIEZO1 activation by Yoda1 induces intracellular Ca^2+^ influx in 2D monolayers, and was associated with the upregulation of p-CaMKII. Moreover, by using p-CaMKII as a marker, we observed the Ca^2+^ influx *in vivo* during decidualization. Previous studies reported that Ca^2+^-CaMK plays an important role in the decidualization process since blocking Ca^2+^ influx with Gd^3+^ or inhibiting CaMK result in impaired decidualization of HESCs [48, 65]. In addition, the decidualization enhanced by Yoda1 is completely abolished by BAPTA-AM and KN93, suggesting that the PIEZO1-mediated decidualization is through its regulatory function as a Ca^2+^ ion channel modulator. Consistent with our data, other studies showed increased intracellular Ca^2+^ upon *in vitro* decidualization of HESCs [65, 66]. However, another study showed that L-type voltage-dependent Ca^2+^ channel (VDCC) mediated Ca^2+^ influx induced by ionophores inhibits cAMP-promoted decidualization [67], suggesting that the level of intracellular Ca^2+^ plays dual roles, by enhancing decidualization at low levels but inhibiting decidualization at high levels, which can explain the phenomenon that low frequency of cycling mechanical stretch enhances decidualization while a high-frequency stretch impairs it [25, 26]. In 1992, spontaneous Ca^2+^ influx was observed in adherent cultured human decidual cells regardless of any external stimulation [68]. We infer that this Ca^2+^ influx may be generated by the contraction and higher expression of PIEZO1 in decidual cells as we report in this study. Other Ca^2+^ mediators have also been reported to contribute to the increase in intracellular Ca^2+^, such as IP_3_ a sensitive Ca^2+^ channel modulator [66], and TRPC1[65] during *in vitro* decidualization of HESCs.

Inhibition of autophagy via the AMPK-mTOR signaling pathway or by knocking down *ATG7* or *ATG5* genes negatively affects the decidualization of human endometrial cells [30, 31]. Similarly, in mice, autophagy inhibitors significantly inhibit artificially induced decidualization [32], and mice lacking autophagy proteins FIP200 and Atg16L1 display embryo implantation and decidualization failure [33, 34]. In addition, the decidualization process is inhibited under microgravity conditions, associated with decreased autophagy flux [36]. These studies indicate that the process of cellular autophagy is highly related to the decidualization of endometrial stromal cells. Furthermore, autophagy is closely related to the cellular tension and the Ca^2+^ influx [37]. The intracellular Ca^2+^ chelation agent BAPTA-AM inhibits autophagy flux [54]. In our study, the absence of PIEZO1 leads to impaired decidualization of mice and human stromal cells along with an inhibited autophagy flux. Detailed investigation revealed a Yoda1/PIEZO1-Ca^2+^/CaMKII-BECN1 cascade, but not mTOR signaling, controlling autophagy and decidualization is, consistent with a previous study which reported that CaMKII promotes autophagy and differentiation by mediating the phosphorylation of BECN1 at Ser90 in Hela cells [55]. This BECN1-dependent autophagy is necessary and sufficient for the PIEZO1-regulated decidualization, as the silencing of *Beclin1* abolished Yoda1-induced decidualization and Tat-BECN1 treatment fully rescued *Piezo1* knocking out caused decidualization failure. As for the decreased mTOR signaling in our *Piezo1^d/d^* mice, we speculate that this is negative feedback caused by the inhibition of autophagy.

The limitation of this study is its relatively weak link to clinical diseases. Many gynecological diseases such as endometriosis, adenomyosis, recurrent pregnancy loss, and preeclampsia are associated with decidualization impairment or failure. Herein, we demonstrate that PIEZO1 mediated decidualization through autophagy is consistent between mice and human cells. We also report that the expression of *PIEZO1* is significantly decreased in the endometrium of baboons with induced endometriosis, suggesting a possibility that PIEZO1 may contribute to the decidualization impairment in endometriosis. However, we observed a rapid increase in *Piezo1* mRNA expression in response to the stretch in mice, which implies variation among samples collected under different mechanical conditions, providing difficulties for comparing *PIEZO1* levels in human biopsies.

In summary, in this study, we have provided evidence for the important roles of the mechanical microenvironment in regulating endometrial decidualization mediated by its receptor PIEZO1 through Ca^2+^/CaMKII and autophagy cascade in both mice and humans. Our findings have uncovered a novel mechanism that regulates decidualization.

## Methods

### Sex as a biological variable

Our study exclusively examined female mice because the disease modeled is only relevant in females.

### Animals

*Pgr^Cre/+^* mice were kindly provided by Dr. FJ DeMayo and Dr. JP Lydon, *Piezo1^f/f^* (Strain #029213) and *Piezo1^tdTomato^* (Strain #029213) mice were obtained from The Jackson Laboratory. The mice were housed in SPF-level facility of experimental animal center of South China agricultural University, with a cycle of 12 hours light (07:00-19:00) and 12 hours dark (19:00-07:00), allowing free access to water and food. All experimental procedures adhered to the guidelines established by the Institutional Animal Care and Use Committee (IACUC) of South China Agricultural University.

The *Pgr^Cre/+^* mice were mated with *Piezo1^f/f^* mice to generate uterine specific *Piezo1* knockout *Pgr^Cre/+^Piezo1^f/f^*(*Piezo1^d/d^*) mice, and the *Pgr^+/+^Piezo1^f/f^*(*Piezo1^f/f^*) mice were used as control. 6 to 8-week-old females were caged with fertile wildtype male to induce pregnancy, the next day with vaginal plug observation was counted as 0.5 dpc. Uteri from *Piezo1^d/d^*and *Piezo1^f/f^* mice at various stages of pregnancy (0.5-7.5 dpc), along with developing fetuses and placentas at 11.5 dpc and 14.5 dpc, were collected, and histomorphological images were obtained.

Artificial decidualization model was performed as described in [40]. Briefly, ovariectomized 6-week-old mice were allowed to recover for 2 weeks, and then subcutaneously injected with 0.1 mL of E_2_ (1 μg/mL) daily at 9:00 a.m. for three consecutive days (Days 1-3), followed by a 2-day rest. On Day 6, at 8:00 a.m., a progesterone-containing silicone tube (1 cm, 250 mg/mL P_4_, Dow Corning) was implanted subcutaneously. At the same time, mice were subcutaneously injected with 0.1 mL of E_2_ (67 ng/mL) daily for three consecutive days. On Day 8, at 3:00 p.m. (6 hours after the final E_2_ injection), artificial decidualization was induced by scratching the antimesometrial side of one uterine horn six times to induce decidualization, the contralateral horn serving as control. Uterine tissues were collected at 6 hours, 1 day, 2 days, and 4 days post-decidualization for further analysis.

All experimental procedures regarding baboons were approved by the Institutional Animal Care and Use Committee (IACUC) of the Michigan State University. Endometriosis was experimentally induced in adult female baboons (Papio anubis) by i.p. inoculation with menstrual endometrium on two consecutive menstrual cycles, as previously described [69]. This model allows us to study the progression of endometriosis by collecting EUE from these animals at various time points following the induction of the disease.

### Measurement of Serum E2 and P4 Levels

The mouse serum collected was stored at -80°C. The estrogen and progesterone concentration in the mouse serum samples was measured by Shanghai Yanhui Biotechnology Co., Ltd.

### Histology and immunostaining

Tissues fixed in 4% paraformaldehyde (PFA) were subjected to dehydration and embedded in paraffin. The paraffin-embedded tissues were sectioned to a thickness of 5 μm. The sections were dewaxed using xylene, rehydrated with a gradient of alcohol, stained with eosin and hematoxylin, dehydrated again and finally mounted with neutral gum.

Immunostaining was conducted according to the previously described protocol [70]. Briefly, paraffin sections (5 μm) were deparaffinized, rehydrated, and subjected to antigen retrieval by boiling in 10 mM citrate buffer for 10 minutes. Endogenous horseradish peroxidase (HRP) activity was inhibited using a 3% H2O2 solution in methanol. After washing three times with PBS, the sections were incubated at 37°C for 1 hour in 10% horse serum for blocking, followed by overnight incubation with each primary antibody at 4°C. The primary antibodies utilized in this study were anti-PIEZO1 (ab128245, Abcam, Cambridge, UK), anti-RFP (600401376, Rockland), anti-pCaMKII (af3493, Affinit), anti-pmTOR (2971s, Cell Signaling Technology), anti-CDK4 (12790T, Cell Signaling Technology), anti-Cyclin D1 (2978T, Cell Signaling Technology), anti-pERK1/2 (4370, Cell Signaling Technology), anti-tERK1/2 (4695, Cell Signaling Technology). After washing, the sections were incubated with biotinylated rabbit anti-goat IgG antibody (1:200, Zhongshan Golden Bridge, Beijing, China) and streptavidin-HRP complex (1:200, Zhongshan Golden Bridge). Positive signals were visualized using the DAB Horseradish Peroxidase Color Development Kit (Zhongshan Golden Bridge) according to the manufacturer’s protocol. The nuclei were counterstained with hematoxylin. For immunofluorescence, sections were incubated with secondary antibody Alexa Fluor Goat anti-Rabbit 488 (Invitrogen, A11008). DAPI was used for counterstaining for immunofluorescence, and Images were captured using a confocal microscope (Leica, TCS SP8, Germany).

### Primary culture and treatment of mESCs

Endometrial stromal cells were isolated from the uteri of mice at 3.5 dpc. The uteri were longitudinally sectioned, rinsed in Hanks’ balanced salt solution (HBSS), and incubated with 1% (w/v) trypsin and 6 mg/ml dispase in 3.5 mL HBSS for 1 hour at 4 °C, followed by 1 hour at room temperature and 10 minutes at 37 °C. The uterine tissues were then washed with HBSS to remove epithelium and subsequently incubated in 6 mL of HBSS containing 0.15 mg/ml Collagenase I (Invitrogen, 17100–017) at 37 °C for 35 minutes. Primary endometrial stromal cells were cultured in DMEM/F12 medium supplemented with 10% fetal bovine serum (FBS). Primary endometrial stromal cells were treated with 10 nM of E_2_ and 1 μM of P_4_ in DMEM/F12 medium containing 2% charcoal-treated fetal bovine serum (cFBS, Biological Industries) to induce *in vitro* decidualization for 48 hours. Stromal cells were treated with Yoda1 (800nM, HY-18723, MedChemExpress) for 48-hour during decidualization period. Inhibitors and recombinant protein, including BAPTA-AM (10 μM, S7534, Selleck, Shanghai, China), KN93 (5 μM, S6787, Selleck, Shanghai, China), U0126 (25 μM, S1102, Selleck, Shanghai, China), and Tat-BECLIN1 (2 μM, S8595, Selleck, Shanghai, China), were applied 1 hour prior to Yoda1, respectively.

### Culture and treatment of HESCs

Human endometrial stromal cells (ATCC, CRL-4003TM) were seeded in 12-well plates and cultured in DMEM/F12 supplemented with 10% charcoal-treated fetal bovine serum (cFBS, Biological Industries) until approximately 80% confluence at 37 °C and 5% CO_2_. *in vitro* decidualization was performed as previously described [40]. Briefly, HESCs were induced with 500 μM dibutyryl cyclic adenosine monophosphate (db-cAMP, D0627, Merck) and 1 μM medroxyprogesterone acetate (MPA, M1629, Merck) for the indicated durations. The treatment with Yoda1 and all inhibitors was consistent with those used for mESCs.

### RNA silencing

The siRNAs targeting human *PIEZO1* (*siPIEZO1*, 5′-GGGACTGCCTCATTCTGTA-3′) and mouse *Beclin1* (*siBeclin1*, 5′-GGCACAATCAATAATTTCA-3′) were designed and synthesized by Ribobio Co., Ltd. (Guangzhou, China). Following the manufacturer’s protocol, mESCs or HESCs were transfected with each *hPIEZO1* siRNA or <u>siBeclin1</u> siRNA using Lipofectamine 2000 Transfection Reagent (Invitrogen, Grand Island, NY) for 6 hours and 24 hours, respectively. A scramble sequence (siScra, siN0000001-1-5, Ribobio) was used as negative control. Each experiment was repeated at least three times.

### Gel-based cell contraction assay

The gel-based cell contraction assay was performed following the manufacturer’s instructions (Cell Biolabs, CBA-201). Briefly, mESCs were separated and counted prior to the experiment. Subsequently, 4.77 mL of collagen solution was added to a 10 mL aseptic centrifuge tube, followed by the addition of 1.23 mL of 5× DMEM solution. After mixing, 170 μL of neutralizing reagent was added in four separate increments. The counted cells were taken in volumes ranging from 0.75 mL to 3 mL and mixed immediately. At this stage, the concentration of cells in the mixture is approximately 10^6^ cells/mL. Following the mixing step, 0.5 mL from each sample was added to 24-well plate and incubated for 1 hour, after which the preheated decidualization treatment cocktail was added and cultured for 48 hours. Photographs were then taken, and the area of gel shrinkage was analyzed using Image J software.

### Western Blot

Tissues and cells were lysed using RIPA Lysis Buffer (Yamei, China), and protein concentrations were quantified with the Pierce™ BCA Protein Assay Reagent (Thermo, USA). A total of 10 μg of protein was loaded onto a 10% SDS-PAGE gel and subsequently transferred to a nitrocellulose membrane, blocked for 1 hour, and then incubated with primary antibody overnight at 4 °C. The primary antibodies used in this study included phosphorylated CaMKII (pCaMKII, 1:1000, af3493, Affinit), phosphorylated ERK1/2 (p-ERK1/2, 1:1000, 4370, Cell Signaling Technology), total ERK1/2 (t-ERK1/2, 1:1000, 4695, Cell Signaling Technology), Cyclin D1 (1:1000, 2978, Cell Signaling Technology), CDK4 (1:1000, 23972, Cell Signaling Technology), LC3A/B (1:1000, 4108, Cell Signaling Technology), phosphorylated BECN1 (p-BECN1, 1:1000, PA5-112018, Thermo Fisher Scientific), total BECN1 (t-BECN1, 1:1000, 3495s, Cell Signaling Technology), phosphorylated mTOR (p-mTOR, 1:1000, 2971, Cell Signaling Technology), TUBULIN (1:1000, 2144 S, Cell Signaling Technology), and GAPDH (1:1000, sc-32233, Santa Cruz Biotechnology). After the membranes were incubated with an HRP-conjugated secondary antibody (1:5000, Invitrogen) for 1 hour, the signals were detected using an ECL Chemiluminescent Kit (Millipore, USA).

### Real-time qPCR

Total RNA was isolated using a Trizol RNA reagent (Takara, Berkeley, CA, USA) and reverse-transcribed into cDNA using the HiScript II Reverse Transcriptase kit (Vazyme, Nanjing, China). For real-time PCR, the cDNA was amplified with a ChamQTM Universal SYBR® qPCR Master Mix (Vazyme) on the RotorGene Q system (Bio-Rad, Hercules, CA, USA). Data was analyzed using the 2^-△△Ct method and normalized to the levels of *Rpl19* (mouse) or *GAPDH* (human). The corresponding primer sequences for each gene are provided in Table S1. Each experiment was repeated at least three times.

### Intracellular calcium concentration assay

Fluo-4 AM was obtained from Beyotime (S1060, Shanghai, China) and used according the manufacturer’s instructions. Briefly, the cells were washed three times with preheated HBSS free of penicillin and streptomycin and then incubated in HBSS containing Fluo-4-AM at a ratio of 1:1000 for 40 min. The media was then incubated in 2% CFBS medium for an additional 30 minutes. Finally, cells were fixed with 4% PFA at room temperature for 20 min, and then stained with DAPI. Images were captured using a Confocal microscope (Leica, TCS SP8, Germany), and the fluorescence values were analyzed using ImageJ software.

### Transmission electron microscope imaging

Specimens were prepared for TEM analysis as follows: Cells were fixed in 2.5% glutaraldehyde in 0.1 M phosphate buffer (pH 7.4) at room temperature for 2 hours, then transferred to fresh 2.5% glutaraldehyde buffer overnight at 4°C. On the second day, cells were post-fixed in 1% OsO4 for 2 hours, dehydrated with a graded ethanol series, and embedded in epoxy resin. Ultrathin sections (80 nm) were prepared using an ultramicrotome (Leica EM FC7), placed on a copper grid, and observed under a transmission electron microscope (FEI/Talos L120C).

### Autophagosomes and autolysosomes measurement

The cells were then transduced with LC3-GFP-RFP adenovirus-containing medium (8 × 107 PFU/mL) for 4 hours with 2% cFBS. After the transduction, the medium was replaced with a normal 2% cFBS for 6 hours. Following incubation, the cells were fixed with 4% PFA at room temperature for 20 minutes, and the nuclei were stained with DAPI. Images were captured using a Confocal microscope (Leica, TCS SP8, Germany). Autolysosomes were observed as yellow dots, while autophagosomes appeared as free red dots.

### RNA-Seq and Data Analysis

The Trizol RNA Reagent (Takara, Dalian, China) was utilized to extract total RNA from the decidual uteri. The concentration and integrity of the RNA were measured using the ND-1000 Nanodrop and the Agilent 2100 TapeStation (Novogene Bioinformatic Technology, Beijing, China), respectively. The quality control parameters employed in this study were: A260/A280 ratio ⩾ 1.8, A260/A230 ratio ⩾ 2.0, and RNA integrity number ⩾ 8.0. The TruSeq RNA sample preparation kit (Illumina, San Diego, CA, USA) was utilized to generate cDNA libraries. RNA sequencing was conducted on an Illumina HiSeq 2500 system. Raw data were processed using an in-house computational pipeline. Differentially expressed genes were selected based on the criteria of fold change > 2 and a false discovery rate (FDR) < 0.05. The RNA-seq raw data were deposited in the Gene Expression Omnibus (GEO) under the accession numbers GSE285591 and GSE286131. GO and KEGG analyses were performed using the DAVID online tools. The cutoff for the false discovery rate (FDR) was established at 0.05.

### Statistics

The data were analyzed using GraphPad Prism 8.0. 2-tails Student’s t-test was employed to compare differences between two groups, while comparisons among multiple groups were performed using one-or two-Way ANOVA followed by Tukey test. All experiments were repeated independently at least three times. In the mouse study, each group consisted of at least three mice. Data are presented as mean±SEM unless stated otherwise. p-value of <0.05 was considered significant.

### Study approval

All animal welfare, procedures, and experiments were approved by the Institutional Animal Care and Use Committee (IACUC) of South China Agricultural University, and Michigan State University.

## Author contributions

RWS, KJW, YW, ATF, ZMY, and HLD conceived the study; JWK, YW, YZ, HXL, GYL, YFY, XQH, JYY, CL, RZ, XZL, SSS, YNL, SH, and YS performed experiments; JPL and FJD provide key resources for mice experiments; RWS, JWK, and YW analyzed the data; RWS, JWK, and YW wrote the draft of manuscript; All authors contributed to the reviewing and editing of the manuscript.

## Acknowledgments

The authors thank our colleagues from the laboratory animal center of South China Agricultural University for their excellent support in maintaining animals. This study is funded by the National Natural Science Foundation of China Nos. 31771664, 32471173, and W2433064, the Double First-Class Discipline Promotion Project No. 2023B10564003, and the National Key Research and Development Program of China No. 2018YFC1004400.

## References

1. Gellersen, B. and J.J. Brosens, Cyclic decidualization of the human endometrium in reproductive health and failure. Endocr Rev, 2014. 35(6): p. 851–905.

2. Garrido-Gomez, T., et al., Defective decidualization during and after severe preeclampsia reveals a possible maternal contribution to the etiology. Proc Natl Acad Sci U S A, 2017. 114(40): p. E8468–E8477.

3. Garrido-Gomez, T., et al., Decidualization resistance in the origin of preeclampsia. Am J Obstet Gynecol, 2020.

4. Su, R.W., et al., Decreased Notch pathway signaling in the endometrium of women with endometriosis impairs decidualization. J Clin Endocrinol Metab, 2015. 100(3): p. E433–42.

5. Lucas, E.S., et al., Recurrent pregnancy loss is associated with a pro-senescent decidual response during the peri-implantation window. Commun Biol, 2020. 3(1): p. 37.

6. Ng, S.W., et al., Endometrial Decidualization: The Primary Driver of Pregnancy Health. Int J Mol Sci, 2020. 21(11).

7. Cha, J., X. Sun, and S.K. Dey, Mechanisms of implantation: strategies for successful pregnancy. Nat Med, 2012. 18(12): p. 1754–67.

8. Su, R.W. and A.T. Fazleabas, Implantation and Establishment of Pregnancy in Human and Nonhuman Primates. Adv Anat Embryol Cell Biol, 2015. 216: p. 189–213.

9. Sternberg, A.K., et al., How Mechanical Forces Change the Human Endometrium during the Menstrual Cycle in Preparation for Embryo Implantation. Cells, 2021. 10(8).

10. Di, X., et al., Cellular mechanotransduction in health and diseases: from molecular mechanism to therapeutic targets. Signal Transduct Target Ther, 2023. 8(1): p. 282.

11. Kefauver, J.M., A.B. Ward, and A. Patapoutian, Discoveries in structure and physiology of mechanically activated ion channels. Nature, 2020. 587(7835): p. 567–576.

12. Coste, B., et al., Piezo1 and Piezo2 are essential components of distinct mechanically activated cation channels. Science, 2010. 330(6000): p. 55–60.

13. Yang, X., et al., Structure deformation and curvature sensing of PIEZO1 in lipid membranes. Nature, 2022. 604(7905): p. 377–383.

14. Lin, Y.C., et al., Force-induced conformational changes in PIEZO1. Nature, 2019. 573(7773): p. 230–234.

15. Li, J., et al., Piezo1 integration of vascular architecture with physiological force. Nature, 2014. 515(7526): p. 279–282.

16. Wang, L., et al., Mechanical sensing protein PIEZO1 regulates bone homeostasis via osteoblast-osteoclast crosstalk. Nat Commun, 2020. 11(1): p. 282.

17. Romac, J.M., et al., Piezo1 is a mechanically activated ion channel and mediates pressure induced pancreatitis. Nat Commun, 2018. 9(1): p. 1715.

18. Friedrich, E.E., et al., Endothelial cell Piezo1 mediates pressure-induced lung vascular hyperpermeability via disruption of adherens junctions. Proc Natl Acad Sci U S A, 2019. 116(26): p. 12980–12985.

19. Atcha, H., et al., Mechanically activated ion channel Piezo1 modulates macrophage polarization and stiffness sensing. Nat Commun, 2021. 12(1): p. 3256.

20. Choi, D., et al., Piezo1-Regulated Mechanotransduction Controls Flow-Activated Lymphatic Expansion. Circ Res, 2022. 131(2): p. e2–e21.

21. Solis, A.G., et al., Mechanosensation of cyclical force by PIEZO1 is essential for innate immunity. Nature, 2019. 573(7772): p. 69–74.

22. Baghdadi, M.B., et al., PIEZO-dependent mechanosensing is essential for intestinal stem cell fate decision and maintenance. Science, 2024. 386(6725): p. eadj7615.

23. Everson, R.B., et al., Extraction of DNA from cryopreserved clotted human blood. Biotechniques, 1993. 15(1): p. 18–20.

24. Abbas, Y., et al., Tissue stiffness at the human maternal-fetal interface. Hum Reprod, 2019. 34(10): p. 1999–2008.

25. Harada, M., et al., Mechanical stretch upregulates IGFBP-1 secretion from decidualized endometrial stromal cells. Am J Physiol Endocrinol Metab, 2006. 290(2): p. E268–72.

26. Saito, R., et al., High stretch cycling inhibits the morphological and biological decidual process in human endometrial stromal cells. Reprod Med Biol, 2020. 19(4): p. 378–384.

27. Kim, J., et al., Acquired contractile ability in human endometrial stromal cells by passive loading of cyclic tensile stretch. Sci Rep, 2020. 10(1): p. 9014.

28. Fleming, A., et al., The different autophagy degradation pathways and neurodegeneration. Neuron, 2022.

29. Hansen, M., D.C. Rubinsztein, and D.W. Walker, Autophagy as a promoter of longevity: insights from model organisms. Nat Rev Mol Cell Biol, 2018. 19(9): p. 579–593.

30. Zhang, Y., et al., AMPK/mTOR downregulated autophagy enhances aberrant endometrial decidualization in folate-deficient pregnant mice. J Cell Physiol, 2021. 236(11): p. 7376–7389.

31. Mestre Citrinovitz, A.C., T. Strowitzki, and A. Germeyer, Decreased Autophagy Impairs Decidualization of Human Endometrial Stromal Cells: A Role for ATG Proteins in Endometrial Physiology. Int J Mol Sci, 2019. 20(12).

32. Su, Y., et al., Endometrial autophagy is essential for embryo implantation during early pregnancy. J Mol Med (Berl), 2020. 98(4): p. 555–567.

33. Oestreich, A.K., et al., *The autophagy protein,* FIP200 (RB1CC1) mediates progesterone responses governing uterine receptivity and decidualizationdagger. Biol Reprod, 2020. 102(4): p. 843–851.

34. Oestreich, A.K., et al., The Autophagy Gene Atg16L1 is Necessary for Endometrial Decidualization. Endocrinology, 2020. 161(1).

35. Zhao, X., et al., Deciphering the Mechanism of Bushen Huoxue Decotion on Decidualization by Intervening Autophagy via AMPK/mTOR/ULK1: A Novel Discovery for URSA Treatment. Front Pharmacol, 2022. 13: p. 794938.

36. Cho, H.J., et al., Microgravity inhibits decidualization via decreasing Akt activity and FOXO3a expression in human endometrial stromal cells. Sci Rep, 2019. 9(1): p. 12094.

37. Shim, M.S., et al., Primary cilia and the reciprocal activation of AKT and SMAD2/3 regulate stretch-induced autophagy in trabecular meshwork cells. Proc Natl Acad Sci U S A, 2021. 118(13).

38. Hu, M., et al., Substrate stiffness differentially impacts autophagy of endothelial cells and smooth muscle cells. Bioact Mater, 2021. 6(5): p. 1413–1422.

39. Braundmeier, A.G. and A.T. Fazleabas, The non-human primate model of endometriosis: research and implications for fecundity. Mol Hum Reprod, 2009. 15(10): p. 577–86.

40. Su, R.W., et al., Aberrant activation of canonical Notch1 signaling in the mouse uterus decreases progesterone receptor by hypermethylation and leads to infertility. Proc Natl Acad Sci U S A, 2016. 113(8): p. 2300–5.

41. Strakova, Z., et al., In vivo infusion of interleukin-1beta and chorionic gonadotropin induces endometrial changes that mimic early pregnancy events in the baboon. Endocrinology, 2005. 146(9): p. 4097–104.

42. Oliver, C., et al., Human decidual stromal cells express alpha-smooth muscle actin and show ultrastructural similarities with myofibroblasts. Hum Reprod, 1999. 14(6): p. 1599–605.

43. Chen, S.T., et al., Embryo-derive TNF promotes decidualization via fibroblast activation. Elife, 2023. 12.

44. Chaudhuri, O., et al., Effects of extracellular matrix viscoelasticity on cellular behaviour. Nature, 2020. 584(7822): p. 535–546.

45. Racioppi, L. and A.R. Means, Calcium/calmodulin-dependent kinase IV in immune and inflammatory responses: novel routes for an ancient traveller. Trends Immunol, 2008. 29(12): p. 600–7.

46. De Koninck, P. and H. Schulman, Sensitivity of CaM kinase II to the frequency of Ca2+ oscillations. Science, 1998. 279(5348): p. 227–30.

47. Wang, Z., G. Palmer, and L.C. Griffith, Regulation of Drosophila Ca2+/calmodulin-dependent protein kinase II by autophosphorylation analyzed by site-directed mutagenesis. J Neurochem, 1998. 71(1): p. 378–87.

48. Lee, S.Y., et al., Nitration of protein phosphatase 2A increases via Epac1/PLCepsilon/CaMKII/HDAC5/iNOS cascade in human endometrial stromal cell decidualization. FASEB J, 2020. 34(11): p. 14407–14423.

49. Kusama, K., et al., Possible roles of the cAMP-mediators EPAC and RAP1 in decidualization of rat uterus. Reproduction, 2014. 147(6): p. 897–906.

50. Lee, C.H., et al., Extracellular signal-regulated kinase 1/2 signaling pathway is required for endometrial decidualization in mice and human. PLoS One, 2013. 8(9): p. e75282.

51. Kandilci, A. and G.C. Grosveld, SET-induced calcium signaling and MAPK/ERK pathway activation mediate dendritic cell-like differentiation of U937 cells. Leukemia, 2005. 19(8): p. 1439–45.

52. Kupzig, S., S.A. Walker, and P.J. Cullen, The frequencies of calcium oscillations are optimized for efficient calcium-mediated activation of Ras and the ERK/MAPK cascade. Proc Natl Acad Sci U S A, 2005. 102(21): p. 7577–82.

53. Illario, M., et al., Calcium/calmodulin-dependent protein kinase II binds to Raf-1 and modulates integrin-stimulated ERK activation. J Biol Chem, 2003. 278(46): p. 45101–8.

54. Bootman, M.D., et al., The regulation of autophagy by calcium signals: Do we have a consensus? Cell Calcium, 2018. 70: p. 32–46.

55. Li, X., et al., CaMKII-mediated Beclin 1 phosphorylation regulates autophagy that promotes degradation of Id and neuroblastoma cell differentiation. Nat Commun, 2017. 8(1): p. 1159.

56. Kang, R., et al., The Beclin 1 network regulates autophagy and apoptosis. Cell Death Differ, 2011. 18(4): p. 571–80.

57. Tang, D., et al., Endogenous HMGB1 regulates autophagy. J Cell Biol, 2010. 190(5): p. 881–92.

58. Wang, J., et al., A non-canonical MEK/ERK signaling pathway regulates autophagy via regulating Beclin 1. J Biol Chem, 2009. 284(32): p. 21412–24.

59. Shoji-Kawata, S., et al., Identification of a candidate therapeutic autophagy-inducing peptide. Nature, 2013. 494(7436): p. 201–6.

60. Vining, K.H. and D.J. Mooney, Mechanical forces direct stem cell behaviour in development and regeneration. Nat Rev Mol Cell Biol, 2017. 18(12): p. 728–742.

61. Han, Y.L., et al., Cell contraction induces long-ranged stress stiffening in the extracellular matrix. Proc Natl Acad Sci U S A, 2018. 115(16): p. 4075–4080.

62. Gnecco, J.S., et al., Organoid co-culture model of the human endometrium in a fully synthetic extracellular matrix enables the study of epithelial-stromal crosstalk. Med, 2023. 4(8): p. 554–579 e9.

63. Gnecco, J.S., et al., Hemodynamic forces enhance decidualization via endothelial-derived prostaglandin E2 and prostacyclin in a microfluidic model of the human endometrium. Hum Reprod, 2019. 34(4): p. 702–714.

64. Xiao, B., Levering Mechanically Activated Piezo Channels for Potential Pharmacological Intervention. Annu Rev Pharmacol Toxicol, 2020. 60: p. 195–218.

65. Kawarabayashi, Y., et al., Critical role of TRPC1-mediated Ca(2)(+) entry in decidualization of human endometrial stromal cells. Mol Endocrinol, 2012. 26(5): p. 846–58.

66. Sohn, J.O., et al., Alterations in intracellular Ca(2+) levels in human endometrial stromal cells after decidualization. Biochem Biophys Res Commun, 2019. 515(2): p. 318–324.

67. Kusama, K., et al., Regulatory Action of Calcium Ion on Cyclic AMP-Enhanced Expression of Implantation-Related Factors in Human Endometrial Cells. PLoS One, 2015. 10(7): p. e0132017.

68. Sartor, P., et al., Voltage-dependent calcium current in human decidual cells and its relation to prolactin secretion. Endocrinology, 1992. 130(6): p. 3433–40.

69. Joshi, N.R., et al., Altered expression of microRNA-451 in eutopic endometrium of baboons (Papio anubis) with endometriosis. Hum Reprod, 2015. 30(12): p. 2881–91.

70. Kang, J.W., et al., Aberrant activated Notch1 promotes prostate enlargement driven by androgen signaling via disrupting mitochondrial function in mouse. Cell Mol Life Sci, 2024. 81(1): p. 155.

